# Striatal acetylcholine enables latent-state creation during reversal learning

**DOI:** 10.64898/2026.07.29.741321

**Authors:** Danielle C. Lowes, Deniz Yagmur Urey, Sofia O. Fernandez, Ines F. Aitsahalia, Samantha R. Ennis, Marco A.M. Prado, Kiyohito Iigaya, Christoph Kellendonk

## Abstract

Like dopamine, acetylcholine is modulated in the striatum by reward-predicting cues and outcomes, yet its computational role remains unclear. Here, we manipulated dorsomedial striatal acetylcholine in mice performing a reversal learning task using genetic and physiologically guided optogenetic approaches. Inhibiting phasic acetylcholine modulation impaired reversal learning, while enhancing modulation during non-rewarded trials facilitated reversal learning. Trial-by-trial acetylcholine dynamics were best explained by a reinforcement learning model in which new latent states are created when experience is poorly explained by existing states. A circuit-constrained model further suggested that acetylcholine promotes plasticity when reward-omission is ambiguous, leading to a surprising prediction: making reward omission explicit should reduce the need for acetylcholine. We confirmed this in a new experiment in which an auditory omission cue substantially reduced the reversal-learning deficit caused by genetic acetylcholine knockdown. These findings suggest that striatal acetylcholine supports reversal learning by promoting state construction in reinforcement learning.

## Introduction

Acetylcholine (ACh) is a neurotransmitter that modulates neuronal activity via muscarinic and nicotinic receptors. Within the striatum ACh is mostly released from cholinergic interneurons (CINs), which comprise 1-2% of striatal neurons with the remaining ACh originating from long-range midbrain projections^1,2^. Classical electrophysiological studies in non-human primates found that presumed cholinergic interneurons modulate their activity in response to salient cues and outcomes coincident with cue- and outcome-induced changes in dopamine (DA) neuron activity or DA levels^3–8^. These findings in non-human primates have been widely replicated in rodents when measuring CIN neuronal activity with *in vivo* physiology or task-evoked changes in ACh using the recently developed genetically encoded sensors for ACh^9–16^.

For DA there is a widely established conceptual framework suggesting that DA acts as several forms of prediction errors^17–22^ that are predicted by reinforcement learning models (though see:^23^). These prediction errors differ in striatal subregions with reward prediction error observed mostly in ventral striatum and state, sensory, and action prediction errors observed in subregions of the dorsal striatum^24–26^. In contrast, the computational role for ACh is still unknown. Some studies observed that changes in cholinergic activity and ACh levels at reward outcome are sensitive to reward expectation^6,7,10^ and possibly may encode a reward prediction error^27^. However, other studies do not support this relationship^5,15^. One computationally-derived concept suggests that striatal ACh is modulated by uncertainty^28^, although this has not been directly supported by empirical evidence.

In this manuscript, we focused on the dorsomedial striatum (DMS) where ACh has been shown to regulate behavioral flexibility including reversal learning^29–34^. Reversal learning is a form of cognitive flexibility impaired in a variety of psychiatric and neurological disorders^35^. This behavior requires a range of brain regions, such as the prefrontal^36^ and orbitofrontal^37^ cortices and the dorsal striatum^38^. In the DMS, extracellular ACh release increases at the onset of a contingency reversal^34^. Inhibition of dorsomedial CINs prevents this release and impairs reversal learning, as does genetic ablation of DMS CINs^32,33^.

Here, we measured changes in ACh and DA levels during a reversal learning task with probabilistic reward. In this task, the mice first learned stable reward contingencies over 5 days, followed by an unexpected reversal in the outcome probabilities. We found that ACh was modulated by both outcome and outcome expectation, with the strongest modulation in non-rewarded trials when reward is expected. Using a conditional genetic knockdown of ACh release and optogenetics, we determined that attenuating phasic ACh release impairs reversal learning while enhancing the ACh modulation in non-rewarded trials facilitates reversal learning. Surprisingly, these manipulations did not affect DA release.

We used computational modeling to identify ACh’s role in reversal learning. Standard reinforcement learning models assume a predefined latent state-space, over which values are updated using reward prediction errors^39^. But naïve animals must construct task-relevant states from experience, and how they do so remains poorly understood^40–42^. Our computational modeling suggested that trial-by-trial ACh dynamics were best explained by a model-derived probability of creating a new latent state, reflecting the mismatch between current experience and past latent states. A circuit-constrained model suggested that ACh-dependent gain enables corticostriatal plasticity when reward omission is ambiguous because it lacks an explicit sensory cue. This model made a testable prediction: externally cueing reward omission should reduce the need for ACh. We performed an additional experiment and confirmed this prediction: an auditory cue at reward omission substantially reduced the reversal-learning deficit caused by genetic knockdown of ACh release.

## Results

### Striatal ACh signals reward outcome and is modulated by expectation

To study reversal learning behavior, we trained food-restricted mice on a probabilistic reversal learning task. Briefly, mice were presented with the option of two levers, one with a 20% chance of yielding a reward, the low probability of reward (P_reward_) lever, and the other with an 80% chance of yielding a reward, the high P_reward_ lever. After five days of acquisition sessions in this task (Acquisition), the reward probabilities were switched once and followed by 5 days with the reversed rule (Reversal). During Reversal the previous low probability lever had an 80% chance of yielding a reward and the previous high probability lever had a 20% chance of yielding a reward (**Figure 1A**). To investigate how ACh and DA release in the dorsomedial striatum contribute to reversal learning, we expressed the fluorescent biosensors GRAB_ACh3.0_ (GACh) and dLight1.3b (dLight) in separate hemispheres for simultaneous imaging with fiber photometry (**Figure 1B**). Mice learned to prefer the high P_reward_ lever during Acquisition and learned to switch to the new high P_reward_ lever during Reversal (**Figure 1C, D**), as evidenced by a decrease in perseverative error streak duration (**Figure 1E**).

**Figure 1:**
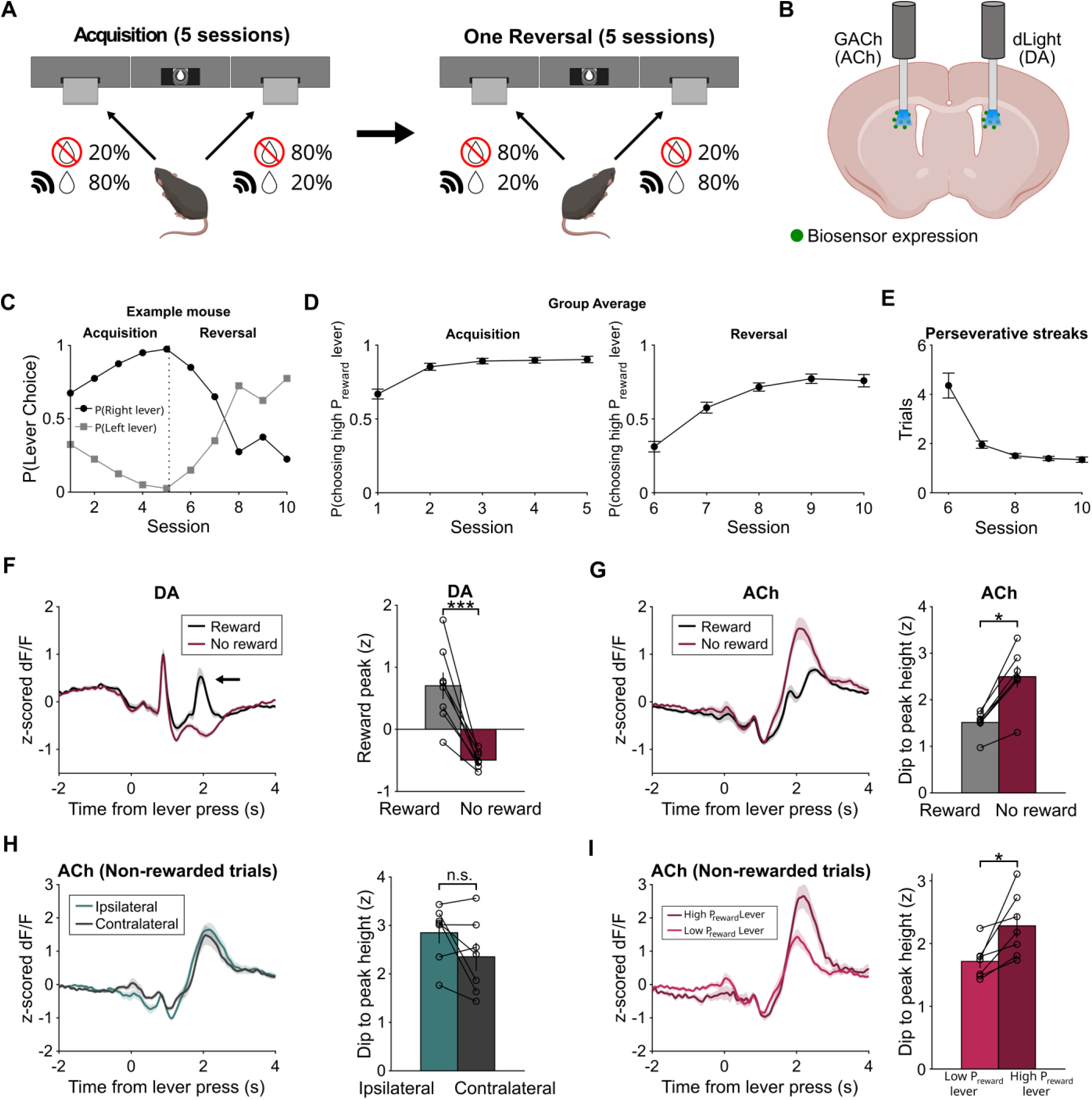
Acetylcholine in the dorsomedial striatum is modulated by reward outcome and expectancy. A) Probabilistic reversal learning task. Mice learned to lever press in an operant task where one lever has an 80% probability of yielding a reward (cued by a motor sound) and a 20% chance of no reward (uncued). The other lever has opposite probabilities. After 5 sessions of this task (Acquisition), mice experience a single reversal where the reward probabilities are switched for an additional 5 sessions (Reversal). B) Experimental approach. The fluorescent biosensors GACh (for ACh) and dLight (for DA) were expressed in separate hemispheres of the dorsomedial striatum for fiber photometry recordings. C) Behavior of an example mouse in the probabilistic choice task. For this mouse, the right lever was the high probability of reward (P_reward_) lever during Acquisition and the low P_reward_ lever during Reversal. D) Group average of performance as measured by choosing the high P_reward_ in the probabilistic choice task. N=19 mice. E) The average perseverative streak length (consecutive low P_reward_ lever choices) across Reversal sessions. N=19 mice. F) Left: DA modulation aligned to lever press (0s) during rewarded (black) and non-rewarded (burgundy) trials averaged for all trials. Arrow designates peak of interest. Right: Quantification of the peak value of the DA signal from 1.7-2.3s after lever press. Paired t-test, ***P=4.9e-4. N=8 mice. G) Left: ACh modulation aligned to lever press (0s) during rewarded (black) and non-rewarded (burgundy) trials averaged for all trials. Right: Quantification of the dip to peak value of the ACh signal. Signed-rank test, *P=0.016. N=7 mice. H) Left: ACh modulation during non-rewarded trials where the mouse pressed a lever ipsilateral (teal) or contralateral to its fiber implant (gray). Right: Quantification of the dip to peak value of the ACh signal. Paired t-test, P=0.12. N=7 mice. I) Left: ACh modulation during non-rewarded trials where the mouse pressed the high P_reward_ lever (burgundy) or the low P_reward_ lever (light burgundy) averaged for Reversal trials. Right: Quantification of the dip to peak value of the ACh signal. Paired t-test, *P=0.015. N=7 mice.

Using fiber photometry, we observed a strong DA transient following the retraction of the lever which produces a noise that serves as a reward-predicting cue (**Figure 1F**). This was followed by a second DA modulation at the time the mouse realizes whether or not the reward dipper has been extended. Rewarded and non-rewarded trials differ in that rewarded trials are cued with the auditory stimulus noise that accompanies dipper extension (**Figure 1A**), while the non-rewarded trials are un-cued and the mouse needs to learn that it is not getting rewarded. Consistent with DA encoding of reward prediction error, fluorescence increased on rewarded trials and decreased on non-rewarded trials (**Figure 1F**).

Like DA, ACh is modulated by reward outcome. GACh fluorescence showed a dip in response to lever retraction which was followed by a rebound in rewarded and non-rewarded trials, with a larger rebound in non-rewarded trials (**Figure 1G**). This difference is not explained by ipsilateral versus contralateral movements which have been shown to modulate ACh in the dorsomedial striatum^9,15^ (**Figure 1H**). In addition to outcome modulation, the ACh signal is modulated by reward expectation, with the largest rebound occurring in non-rewarded trials when the reward is expected (**Figure 1I**).

### Striatal VAChT knockout impairs reversal learning but preserves acquisition

To address a causal role of striatal ACh modulation in reversal learning behavior, we crossed floxed vesicular acetylcholine transporter homozygous mice (VAChT^flx/flx^) with Drd2-Cre heterozygous, floxed VAChT homozygous mice (D2Cre^+/-^-VAChT^flx/flx^) to obtain Drd2-Cre^+/-^-VAChT^flx/flx^ (VAChTcKO) and VAChT^flx/flx^ (control) offspring. This genetic strategy has been shown to greatly reduce baseline and task-evoked ACh release from CINs in the dorsal striatum while sparing VAChT expression in cholinergic neurons of the basal forebrain and midbrain (**Figure 2A**)^43^. Moreover, it does not affect glutamate that is co-released from CINs. Mice were first trained to lever press in a continuous reinforcement schedule (CRF). VAChTcKO mice did not exhibit any deficits in lever press latency or reward retrieval latency, suggesting intact instrumental learning (**Figure S1A, B**). We then used fiber photometry to measure GACh and dLight fluorescence during the probabilistic reversal learning task and confirmed that VAChTcKO mice exhibit blunted ACh release in the dorsomedial striatum (**Figures 2B, C**). Surprisingly, we did not observe any differences in DA release (**Figures 2D, E & S1C**), suggesting limited modulation of DA release by ACh under our task conditions^16,44^. At the behavioral level, we found that VAChTcKO exhibited intact acquisition of the task but impaired reversal learning, as measured by comparing high P_reward_ lever choice per session between groups (**Figure 2F, G**). Furthermore, VAChTcKO mice exhibit longer streaks of perseverative errors during the first three sessions of reversal learning (**Figure 2H**). These results are consistent with a reported deficit in a deterministic reversal learning task^16^ and extend the finding to a probabilistic reversal learning task.

**Figure 2:**
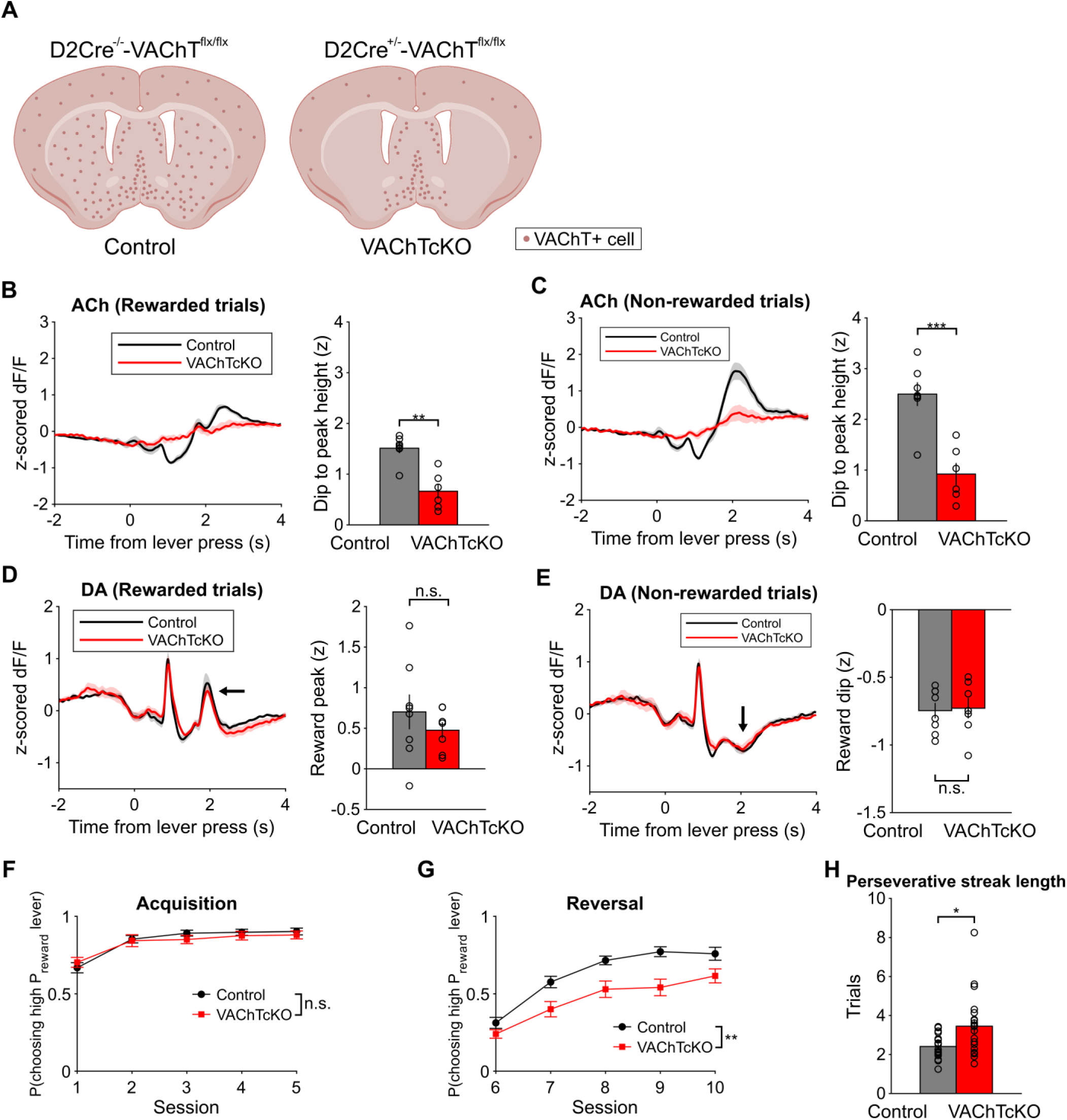
Decreasing acetylcholine release impairs reversal learning. A) Representation of the effect of the VAChTcKO genetic model on VAChT expression, based on Favier et al. 2020. VAChTcKO reduces VAChT mRNA in cholinergic interneurons of the striatum but spares expression in medial septum, basal forebrain and brain stem projections. B) Left: ACh modulation aligned to lever press (0s) during rewarded trials in Control (black) and VAChTcKO (red) mice. Right: Quantification of the dip to peak value of the ACh signal. Rank-sum test, **P=0.0023. N=7 control, N=6 VAChTcKO. C) Left: ACh modulation during non-rewarded trials in Control (black) and VAChTcKO (red) mice. Right: Quantification of the dip to peak value of the ACh signal. Two-sample t-test, ***P=5.1e-4. N=7 control, N=6 VAChTcKO. D) Left: DA modulation during rewarded trials in Control (black) and VAChTcKO (red) mice. Right: Quantification of the peak value of the DA signal from 1.7-2.3s after lever press. Two-sample t-test, P=0.38. N=8 control, N=7 VAChTcKO. E) Left: DA modulation during non-rewarded trials in Control (black) and VAChTcKO (red) mice. Right: Quantification of the minimum value of the DA signal from 1.7-2.3s after lever press. Two-sample t-test, P=0.86. N=8 control, N=7 VAChTcKO. F) Mouse behavior during Acquisition sessions. There was no difference in choosing the high P_reward_ lever between Control (black) and VAChTcKO mice (red). 2-way repeated measures ANOVA, P=0.64 no main effect of genotype, P=0.58 no session*genotype interaction. N=19 control, N=20 VAChTcKO. G) Mouse behavior during Reversal sessions. Control mice chose the high P_reward_ lever more often than VAChTcKO mice (red). Mixed-effects analysis. **P=0.0016 main effect of genotype, P=0.10 no session*genotype interaction. N=19 control, N=20 VAChTcKO. H) Average perseverative streak length (consecutive low P_reward_ choices) during first three Reversal sessions. VAChTcKO mice exhibited longer perseverative streak lengths. Rank-sum test, *P=0.017. N=19 control, N= 20 VAChTcKO.

### Tonic CIN stimulation blunts phasic ACh signaling and impairs reversal learning

The conditional knock out of VAChT is not just restricted to the dorsomedial striatum but also targets some cortical and ventral striatal CINs from development on^43^. To blunt ACh modulation selectively in the adult dorsomedial striatum, we injected ChAT-IRES-Cre mice with Cre-dependent red-shifted channelrhodopsin (AAV-hSyn-DIO-ChrimsonR-tdTomato) or control virus (AAV-hSyn-DIO-mCherry) and performed tonic 10 Hz stimulation of dorsomedial CINs during Acquisition and Reversal phases of the probabilistic reversal learning task (**Figure 3A**). Previous studies have shown that CINs exhibit a pause in firing *ex vivo* following depolarizing current injection possibly due to activation of voltage-gated ion channels^45^. We also observed this phenomenon with optogenetic stimulation of CINs, as a single light pulse or pulse train produces an increase and then decrease in the GACh signal (**Figure S2A**). Tonic stimulation strategy avoids this complication by producing a sustained increase in ACh release that we reasoned should become less sensitive to task evoked modulation. Indeed, tonic stimulation enhanced baseline ACh levels (**Figure S2B**) and when Chrimson mice were stimulated with 10 Hz light pulses during the entire session of the probabilistic reversal learning task, outcome-evoked ACh modulation was blunted in rewarded and non-rewarded trials (**Figure 3B, C**). Like VAChTcKO mice, tonic stimulation did not impact acquisition (**Figure 3D**) but impaired reversal learning performance and increased perseverative streak length (**Figure 3E, F**). These results suggest phasic ACh modulation in the adult dorsomedial striatum is required for efficient reversal learning.

**Figure 3:**
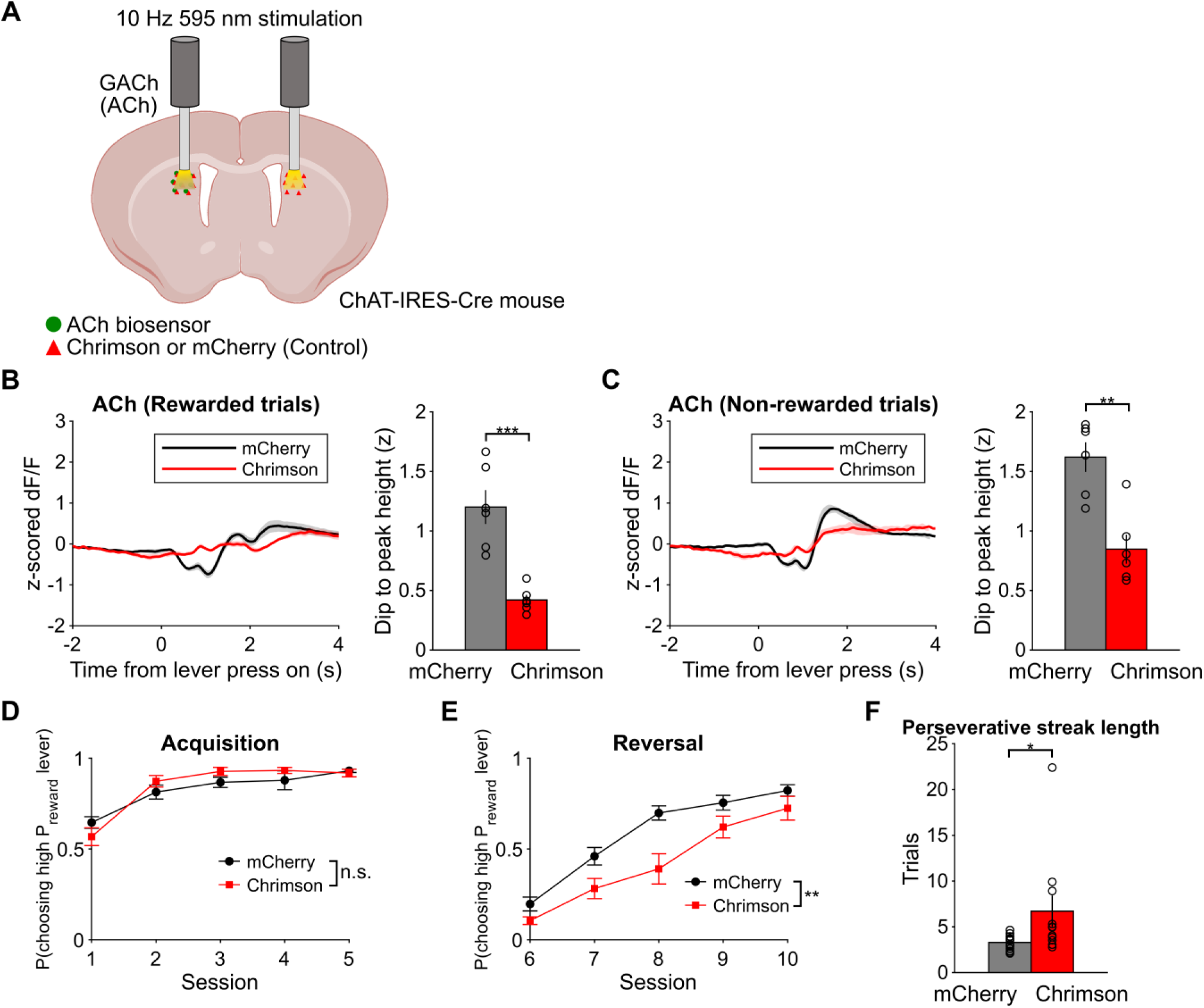
Tonic stimulation of cholinergic interneurons attenuates task evoked changes in acetylcholine levels and impairs reversal learning. A) Experimental schema. ChAT-IRES-Cre mice expressing GACh and Cre-dependent Chrimson or mCherry were given 10 Hz 595 nm stimulation during Acquisition and Reversal sessions. B) Left: ACh modulation during rewarded trials for mCherry control (black) and Chrimson (red) mice. Right: Quantification of the dip to peak value of the ACh signal. Two sample t-test, ***P=3.7e-4. N=6 mCherry, N=6 Chrimson. C) Left: ACh modulation during non-rewarded trials for mCherry control (black) and Chrimson (red) mice. Right: Quantification of the dip to peak value of the ACh signal. Two sample t-test, **P=0.0013. N=6 mCherry, N=6 Chrimson. D) Mouse behavior during Acquisition sessions. There was no difference in choosing the high P_reward_ lever between mCherry control (black) and Chrimson (red) mice. 2-way repeated measures ANOVA. P=0.54 no main effect of virus, P=0.13 no session*virus interaction. N=15 mCherry, N=11 Chrimson. E) Mouse behavior during Reversal sessions. mCherry mice chose the high P_reward_ lever more often than Chrimson mice. Mixed-effects analysis. **P=0.0056 main effect of virus, *P=0.018 session*virus interaction. F) Average perseverative streak length during first three Reversal sessions. Chrimson mice exhibited longer perseverative streak lengths than mCherry mice. Rank-sum test, *P=0.013. N=15 mCherry, N=11 Chrimson.

### Physiologically guided inhibition of DMS CINs enhances phasic ACh signaling and improves reversal learning

The VAChTcKO mice show a deficit perseverating on the previously high P_reward_ lever even if it is not rewarded, and ACh modulation is most pronounced after non-rewarded outcome when reward was expected. If this modulation is important, we hypothesized that selective enhancement of the ACh signal at non-rewarded outcomes should facilitate reversal learning. CINs exhibit rebound excitation in their membrane potential following hyperpolarizing current injection^45^. We sought to exploit this intrinsic mechanism to enhance the biphasic dip-to-peak modulation at non-rewarded outcomes. To this end, we injected ChAT-IRES-Cre mice with Cre-dependent halorhodopsin (AAV-hSyn-DIO-eNpHR-mCherry) or control virus (AAV-hSyn-DIO-mCherry) in addition to GACh and dLight biosensors (**Figure 4A**). Consistent with^45^ brief inhibition of CINs with halorhodopsin induces a dip followed by a rebound in the measured GACh signal (**Figure S3A**). We exploited this characteristic of CINs to enhance the phasic ACh signal during the probabilistic reversal learning task. During Acquisition and Reversal, we delivered a light pulse after non-rewarded lever presses and found that it increases the dip-to-peak height of the phasic ACh response (**Figure 4B**; note that some of the mice were stimulated slightly earlier **Figure S3B**). This manipulation did not affect ACh responses on rewarded trials (**Figure 4C**), and it did not affect DA responses on non-rewarded trials or rewarded trials (**Figure 4D, E**). Enhancing ACh dip-to-peak height in non-rewarded trials did not affect acquisition of the task but facilitated reversal learning (**Figure 4F, G**). eNpHR mice exhibited a decrease in perseverative streak length, that was not significant possibly due to a floor effect (**Figure 4H**).

**Figure 4:**
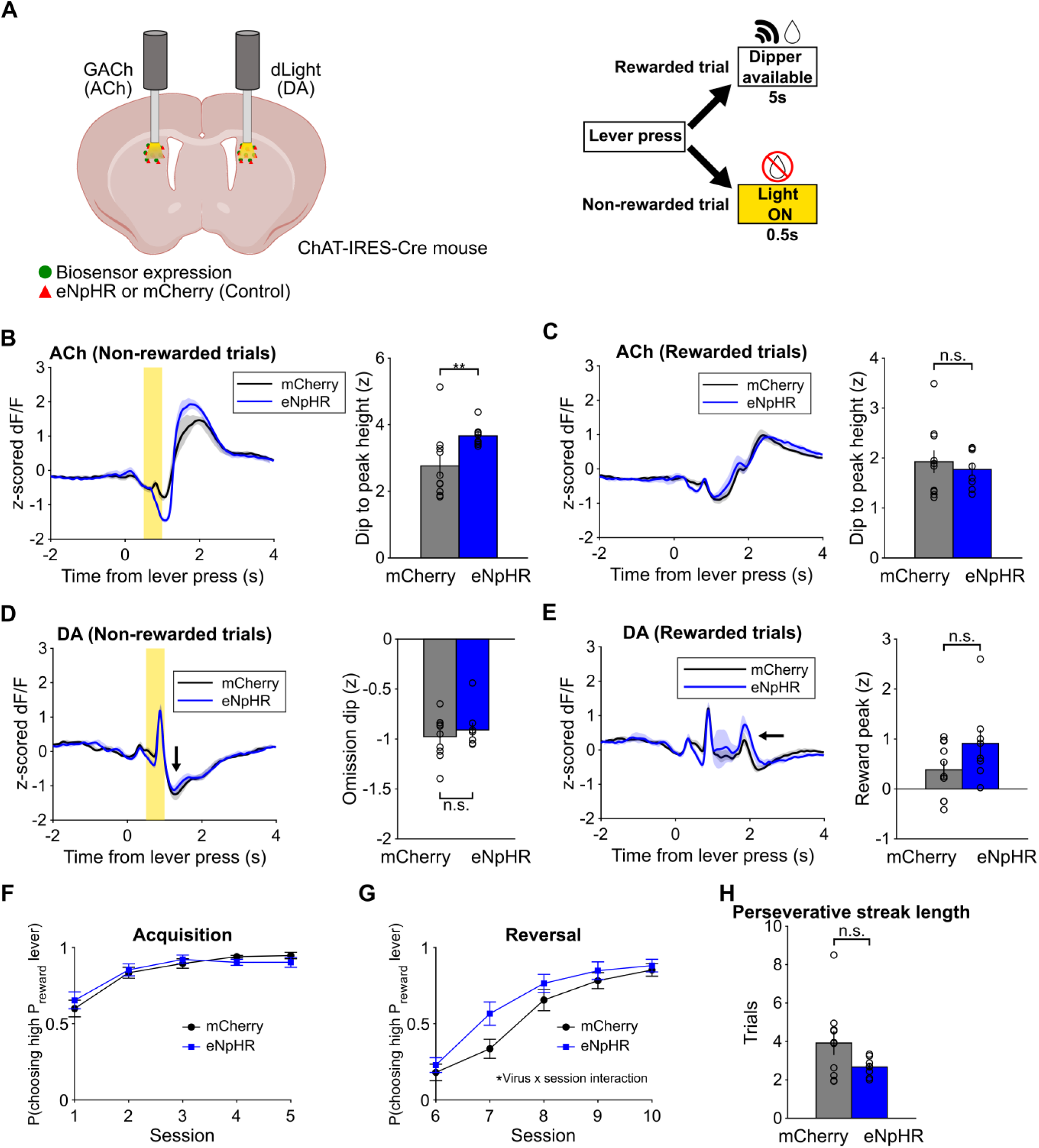
Enhancing acetylcholine modulation at non-rewarded outcome facilitates reversal learning. A) Experimental schema. ChAT-IRES-Cre mice expressing Cre-dependent eNpHR or mCherry were given 500ms pulses of 595 nm light on non-rewarded trials during Acquisition and Reversal sessions. B) Left: ACh modulation during non-rewarded trials for mCherry control (black) and eNpHR (blue) mice. Right: Quantification of the dip to peak value of the ACh signal. Rank-sum test, **P=0.0044. N=10 mCherry, N=8 eNpHR. C) Left: ACh modulation during rewarded trials for mCherry control (black) and eNpHR (blue) mice. Right: Quantification of the dip to peak value of the ACh signal. Two sample t-test, P=0.59. N=10 mCherry, N=8 eNpHR. D) Left: DA modulation during non-rewarded trials in mCherry (black) and eNpHR (blue) mice. Arrow designates dip of interest. Right: Quantification of the minimum value of the DA signal from 1.7-2.3s after lever press. Rank-sum test, P=0.83. N=10 mCherry, N=8 eNpHR. E) Left: DA modulation during rewarded trials in mCherry (black) and eNpHR (blue) mice. Arrow designates peak of interest. Right: Quantification of the peak value of the DA signal from 1.7-2.3s after lever press. Two-sample t-test, P=0.38. N=10 mCherry, N=8 eNpHR. F) Mouse behavior during Acquisition sessions. There was no difference in pressing the high P_reward_ lever between mCherry (black) and eNpHR (blue) mice. 2-way repeated measures ANOVA. P=0.92 no main effect of virus, P=0.39 no session*virus interaction. N=10 mCherry, N=8 eNpHR. G) Mouse behavior during Reversal sessions. eNpHR mice chose the high P_reward_ lever more often than mCherry mice. 2-way repeated measures ANOVA. P=0.18 no main effect of virus, *P=0.049 session*virus interaction. N=10 mCherry, N=8 eNpHR. H) Average perseverative streak length during first three Reversal sessions. Two sample t-test, P=0.10. N=10 mCherry, N=8 eNpHR.

### Computational modeling links ACh to latent-state creation

Our empirical findings showed that ACh in the dorsomedial striatum is necessary for reversal learning and revealed a stereotyped signature: ACh modulation is most pronounced during reward omissions, and selective enhancement of this signal in non-rewarded trials accelerates reversal learning. What computation does this ACh signal carry? Standard reinforcement learning (RL) accounts assume a predefined task state space, but the mice in our task were naïve and had to construct relevant states from experience^42^. We therefore developed an RL model in which the agent creates a new latent state when the current experience cannot be accommodated by the existing states (**Figure 5A**; see Methods for full model specification).

**Figure 5:**
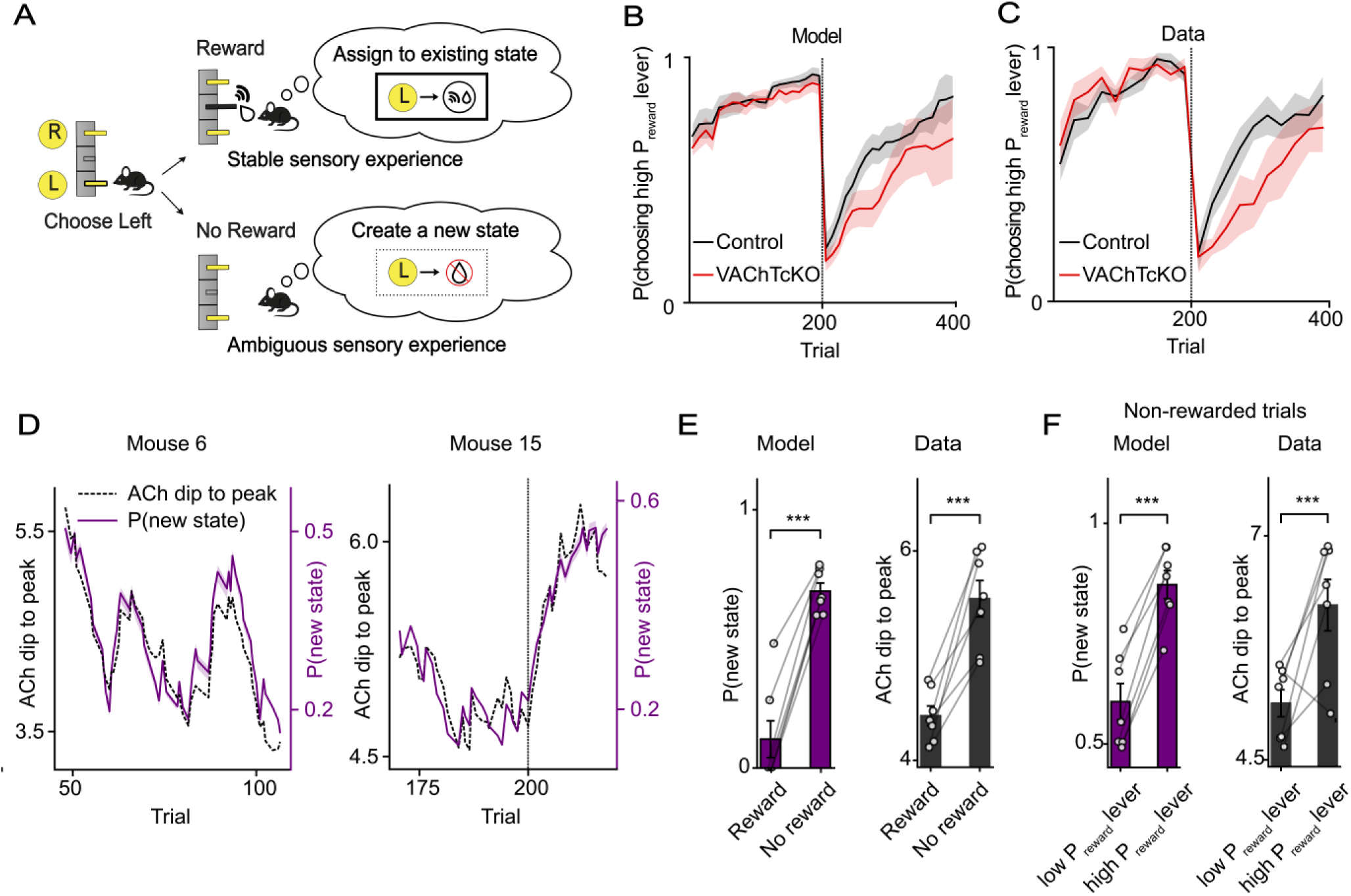
Computational modeling links DMS ACh modulation to latent-state creation. **A)** State-creation reinforcement-learning model. On each trial, the model assigned outcome-related experience either to an existing latent state or to a newly created state. A Chinese Restaurant Process–inspired rule favored assignment to states with similar prior experiences and greater prior use. In this example, the consistent, cue-supported reward-delivery experience was assigned to an existing state (top), whereas the ambiguous, uncued reward-omission experience prompted creation of a new latent state (bottom). **B)** The state-creation reinforcement-learning model captured choice behavior in control and VAChTcKO mice. Learning curves were averaged across simulations from seven control and six VAChTcKO agents. Shading indicates s.e.m. The graphs depict the 200 free choice trials for acquisition and 200 free choice trials for reversal. **C)** Corresponding trial-by-trial choice behavior in seven control and six VAChTcKO mice. Shading indicates s.e.m. **D)** Trial-by-trial state-creation probability predicted DMS ACh modulation. The model was fit to the behavioral data from each control animal and thus the model was blind to the ACh data. The model-derived state-creation probability (purple) captured the observed outcome-locked DMS ACh modulation (black). Two representative control animals are shown. Shading indicates s.e.m. across model simulations. **E)** The model-derived state-creation probability captured the greater DMS ACh modulation on non-rewarded trials during reversal. Both the model (purple) and data from control animals (black) showed significantly greater signals on non-rewarded trials. ***P < 0.001, non-parametric permutation test. Error bars indicate s.e.m. **F)** The model-derived state-creation probability captured the greater DMS ACh modulation following no reward outcomes after selection of the low versus high P_reward_ lever during reversal. Both the model (purple) and data from control animals (black) showed significantly greater signals following selection of the high P_reward_ lever. ***P < 0.001, non-parametric permutation test. Error bars indicate s.e.m.

The model was motivated by Chinese Restaurant Process formulations of latent-cause inference^46^ and state-splitting accounts of extinction learning^47^. On each trial, the model agent selected an action and received an outcome-related experience input. The agent then either assigned this experience to an existing action-specific latent state or created a new state. This decision depended on the similarity of the current input to existing state representations, each defined by the average of inputs previously assigned to that state. Reuse was favored when the current input closely matched an established state and when that state had been used frequently; conversely, a new state was created when the current experience was poorly explained by the best existing state. To reflect continuous-reinforcement pretraining, the model began with an initial reward-delivery state but no pre-existing reward-omission state; additional latent states could then be created for both rewarded and non-rewarded experiences during probabilistic learning. Motivated by the task structure, we represented the reward-delivery experience input as a stable, reproducible pattern that reflects reliable sensory events, such as dipper extension and associated auditory and visual cues. By contrast, reward-omission experience input was represented by a pattern that evolved gradually across encounters because omission carried no corresponding explicit external marker.

The state-creation RL model reproduced the reversal learning trajectory observed in the data (**Figure 5B - model and Figure 5C - data**). In the model with parameters fit to the choice data of control animals, new states were preferentially created following unrewarded outcomes and reversals (**Figure S4**), precisely the conditions under which VAChTcKO mice showed impaired learning. We therefore fit the same state-creation RL model to VAChTcKO choice data and asked whether altered state creation could account for their behavioral deficit. Indeed, the fitted VAChTcKO model showed reduced creation of reward omission-related states (**Figure S5**) and captured key features of VAChTcKO behavior, including impaired reversal learning (**Figures 5B,C**), and prolonged perseveration early after reversal (**Figure S5**). We next asked whether the model-derived latent-state-creation probability could account for DMS ACh dynamics. For each animal, we first quantified the dip-to-peak ACh signal for each individual trial in the experimental data (see Methods). We then used the model fitted to choice data to generate a trial-by-trial prediction of the state-creation probability, P(new state). Although the model was fit only to choice data and not to neural data, the model-derived state-creation probability remarkably predicted trial-by-trial ACh modulation at the level of individual animals (**Figure 5D**).

State creation probability also recapitulated the characteristic outcome-ACh response described in Figure 1, including the larger rebound on non-rewarded trials (**Figure 5E**) and the enhanced response when the high P_reward_ lever was unrewarded (**Figure 5F**). Critically, the model’s trial-by-trial state-creation probability signal was reliably recovered from simulated choices generated by the model and was robust to different model-fitting implementations (**Figure S6**). These results suggest that ACh did not merely track the average task structure; it tracked the animal-specific latent-state creation processes inferred from each animal’s own choices.

We next compared this account with alternative models that could support reversal learning: a standard Q-learning model, an uncertainty-modulated reinforcement learning model in which learning rates are governed by expected and unexpected uncertainty^48,49^; and a hidden meta-state inference model with two predefined contingencies, one with left choice being higher-value than right choice and the other with right choice being higher-value than left choice^50–53^. Whereas the Q-learning provides a standard benchmark, we included the latter two models because those contain uncertainty-related signals that could, in principle, account for ACh modulation^48,49,52^. Behaviorally, these models did not differ significantly from the state-creation model in average cross-validated choice likelihood. This similarity was expected for a simple one-time reversal task, in which most trials occur under the same contingency as the preceding trial, and coarse choice behavior can be captured easily. However, when we simulated the fitted models, the state-creation model more accurately captured the animals’ probability of repeating the same action across trial conditions defined by previous action, previous outcome, reward likelihood, and whether trials occurred before or after reversal (**Figure S7**).

The neural predictions sharply dissociated the models. None of the alternative models accounted for the observed trial-by-trial DMS ACh dynamics (**Figure S8**). In particular, models based on previously proposed uncertainty signals^48,49,52^ did not reproduce the characteristic ACh response pattern conditioned on outcome and reward expectation. Thus, although several models could capture average reversal behavior, only the state-creation RL model jointly accounted for detailed choice structure and trial-by-trial DMS ACh dynamics. Together, these results suggest that DMS ACh is best explained by a behaviorally inferred new state-creation probability.

### A circuit-constrained model links ACh-dependent state creation to corticostriatal plasticity

Our algorithmic model’s analysis offers an interpretable computational function to ACh, but does not specify how such computation could be implemented in striatal circuitry. To address this question, we built a standard circuit-constrained neural network model in which cortical inputs projected to opponent MSN populations within left- and right-action channels of the dorsal striatum^54–56^ (**Figure 6A**). Corticostriatal synapses in the chosen-action MSN channel underwent dopamine-dependent opponent actor plasticity^57^, where corticostriatal weights encoded action-specific outcome experience associations. Because DA release was unaltered in VAChTcKO mice, the model assumed intact dopaminergic teaching signals throughout.

**Figure 6:**
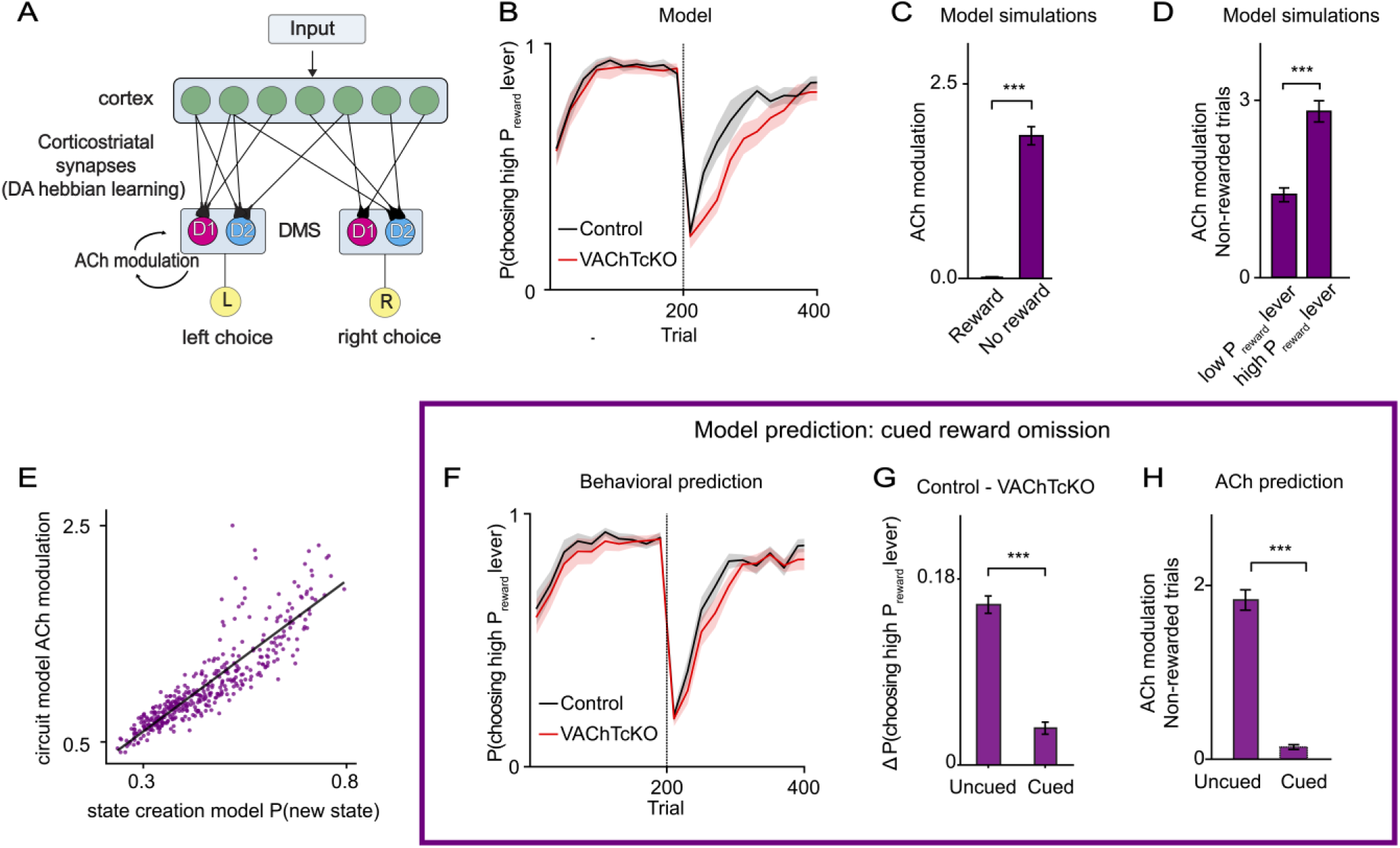
A circuit-constrained model links ACh-dependent state creation to corticostriatal plasticity. **A)** Corticostriatal circuit-constrained model. Outcome-related cortical inputs projected to opponent medium spiny neuron (MSN) populations within left- and right-action channels of the dorsal striatum. Inputs were constructed using the same stable reward and drifting uncued reward-omission structure used in the state-creation reinforcement-learning model. At outcome, a local state–outcome mismatch signal reflecting the mismatch between the plasticity implied by the outcome and the action state supported by existing corticostriatal weights recruited an ACh-dependent increase in outcome-phase gain. This gain enhanced recruitment of outcome representations and dopamine-dependent Hebbian corticostriatal plasticity. **B)** The circuit-constrained model captured choice behavior in control and VAChTcKO mice. Learning curves were averaged across simulations from fifteen control and fifteen VAChTcKO agents. Shading indicates s.e.m. **C)** The circuit model’s mismatch-driven ACh-gain signal captured the greater DMS ACh modulation on non-rewarded trials during reversal. The model, like the data, showed significantly greater signals on non-rewarded trials. ***P < 0.001, non-parametric permutation test. Error bars indicate s.e.m. **D)** The circuit model’s mismatch-driven ACh-gain signal captured the greater DMS ACh modulation following non-rewarded outcomes after selection of the low versus high P_reward_ lever during reversal. The models, like the data, showed significantly greater signals following selection of the high P_reward_ lever. ***P < 0.001, non-parametric permutation test. Error bars indicate s.e.m. **E)** The circuit model’s mismatch-driven ACh-gain signal was consistent with the state-creation probability derived from the state-creation reinforcement-learning model. The circuit model generated synthetic choice data, which were fit with the state-creation model. Across the trial-by-trial mean trajectories, the circuit model’s ACh-dependent gain signal was strongly correlated with state-creation probability derived from the fitted state-creation model (Pearson’s r = 0.88, p <0.0001, Fourier phase randomization test^58^). **F)** The model predicted improved reversal learning in VAChTcKO mice when reward omission was paired with an explicit cue. A reliable cortical representation of reward omission was predicted to reduce the need for ACh-dependent outcome gain and rescue reversal learning in VAChTcKO mice. **G)** Behavioral prediction. The model predicted that the performance difference between control and VAChTcKO mice would be reduced when reward omission was explicitly cued, compared with the original uncued task. ***P < 0.001, non-parametric permutation test. Error bars indicate session-level bootstrap 95% confidence intervals. **H)** ACh prediction. The model predicted that DMS ACh modulation on non-rewarded trials for the control group during reversal would be reduced when reward omission was explicitly cued relative to the original uncued task. ***P < 0.001, non-parametric permutation test. Error bars indicate s.e.m.

Following the reward/no-reward asymmetry introduced above, we instantiated reward and no-reward events differentially at the cortical level. The circuit model used the same mathematical input construction as the state-creation model. The cued reward outcome was modeled as a stable cortical input reflecting reliable sensory events at reward delivery; uncued reward omission was modeled as a weaker, drifting input. Reward, therefore, repeatedly recruited the same corticostriatal ensemble, whereas no reward recruited partially overlapping, shifting ensembles across trials. This asymmetry created a specific challenge for reversal: Conventional corticostriatal plasticity could readily reinforce stable reward-associated ensembles, but weakly reproducible omission-related ensembles were less likely to be recruited reliably into the plasticity update. This impairment is expressed behaviorally as perseveration on the previously rewarded lever.

We then asked what neuromodulatory signal could overcome this failure. The model identified a corticostriatal state–outcome mismatch signal (see Methods for details): a quantity that measures the mismatch between the plasticity implied by outcome and the action state currently supported by corticostriatal weights. This mismatch was largest when existing corticostriatal weights supported an action tendency opposite to that implied by the outcome, for example, when no reward occurred despite the striatum channel promoting that action. We modeled dorsal striatal ACh as an outcome-phase gain-modulation driven by this mismatch. Elevated ACh modulation increased striatal recruitment to plasticity during the outcome period, allowing weak, drifting no-reward experience inputs from the cortex to drive stable, action-specific corticostriatal updates.

The circuit model recapitulated each of our experimental manipulations. Intact ACh gain modulation produced control-like reversal learning (**Figure 6B**). Suppressing ACh gain modulation reproduced the VAChTcKO and tonic stimulation deficits (**Figure 6B**), sparing acquisition but impairing reversal, because uncued reward omissions failed to recruit sufficient MSN activity to update the previously rewarded action. Simulated ACh modulation (DMS gain modulation driven by state-outcome mismatch) also reproduced the lever- and outcome-conditioned dynamics observed in DMS (**Figure 6C, D**).

We found that the circuit model’s state–outcome mismatch signal is closely aligned with the algorithmic model’s state creation signal (**Figure 6E**). To test this, we generated synthetic choices from the circuit model, then fit the state creation reinforcement learning introduced in the previous section to the synthetic choice data. We found that the new state creation probability from the state-creation model matched the circuit-model’s ACh modulation signal across trials. This association remained significant under a Fourier phase-randomization test that preserved the slow temporal structure of the signals^58^ (See Methods), indicating that it was not explained by shared autocorrelation^59^. Thus, corticostriatal state–outcome mismatch provides a candidate circuit implementation of state creation within our model. One modeling interpretation is that when existing experience-action representations stored in corticostriatal activity fail to align with the obtained outcome, this mismatch signal recruits ACh-dependent plasticity to create a new experience-action representation.

The model makes a specific prediction. If ACh is required specifically because uncued reward omission provides an unstable cortical drive, then supplying an explicit no-reward cue (e.g., a tone) should stabilize the cortical representation and reduce the need for ACh-mediated gain. In the model, adding a fixed no-reward cue restored reversal performance in simulated low-ACh agents (**Figure 6F, G**) and reduced simulated ACh modulation for the control group at non-rewarded outcomes (**Figure 6H**). We tested this prediction directly in a new experiment described below.

### Cueing non-rewarded trials reduces reversal learning deficits in VAChTcKO mice

To test the computational model’s prediction that providing a reliable sensory cue for reward omission would reduce the requirement for DMS ACh in reversal learning, we performed a modified version of the probabilistic reversal learning task on VAChTcKO and control mice while recording dorsomedial GACh and dLight signals. In this version, an auditory cue was presented immediately after lever press on non-rewarded trials (**Figure 7A**). As in the original non-cued experiment, task-evoked ACh modulation was markedly blunted in the VAChTcKO mice (**Figure 7B, C**), and acquisition behavior did not differ between genotypes (**Figure 7D**).

**Figure 7:**
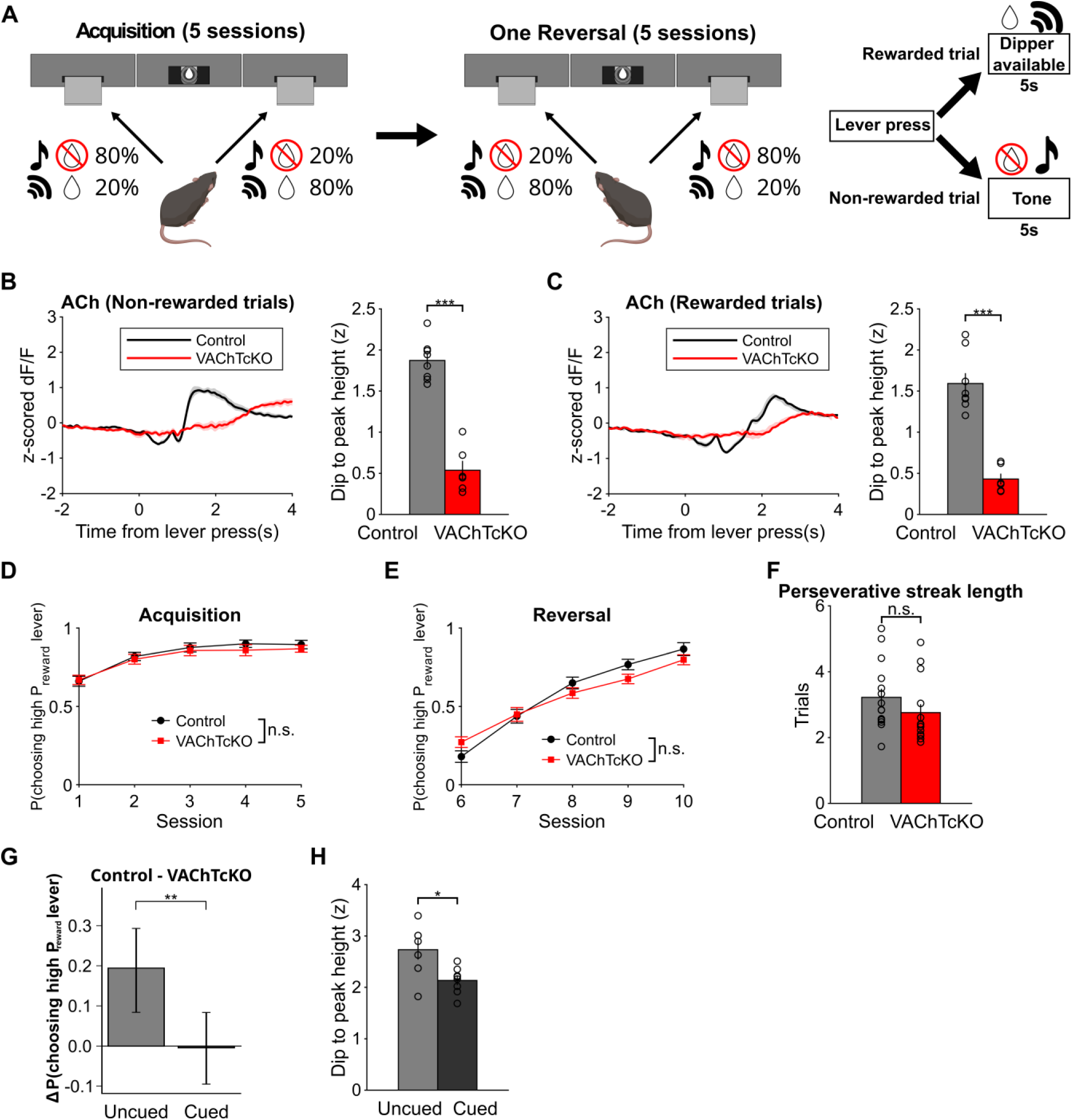
Cueing non-rewarded outcome reduces the reversal learning deficit in VAChTcKO mice. A) Schema of probabilistic reversal learning task as in Figure 1 with the exception that non-rewarded outcomes are now cued by a tone. B) Left: ACh modulation aligned to lever press (0s) during rewarded trials in Control (black) and VAChTcKO (red) mice. Right: Quantification of the dip to peak value of the ACh signal. Two-sample t-test, ***P=6.0e-7. N=8 Control, N=6 VAChTcKO. C) Left: ACh modulation during non-rewarded trials in Control (black) and VAChTcKO (red) mice. Right: Quantification of the dip to peak value of the ACh signal. Two-sample t-test, ***P=8.7e-6. N=8 Control, N=6 VAChTcKO. D) Mouse behavior during Acquisition sessions. There was no difference in choosing the high P_reward_ lever between Control (black) and VAChTcKO mice (red). 2-way repeated measures ANOVA. P=0.53 no main effect of genotype, P=0.85 no session*genotype interaction. N=15 Control, N=15 VAChTcKO. E) Mouse behavior during Reversal sessions. There is no main effect of genotype in choosing the high P_reward_ lever in control (black) versus VAChTcKO mice (red). 2-way repeated measures ANOVA. P=0.51 no main effect of genotype, *P=0.019 session*genotype interaction. N=15 Control, N=15 VAChTcKO. F) Average perseverative streak length (consecutive low P_reward_ choices) during first three Reversal sessions. Perseverative streak length did not differ significantly between genotypes. Rank-sum test, P=0.058. N=15 Control, N=15 VAChTcKO. G) The difference in choosing the high P_reward_ lever between control and VAChTcKO mice during the first three Reversal sessions was significantly reduced in the cued no-reward experiment than in the original non-cued experiment. **P < 0.01, non-parametric permutation test. Error bars indicate session-level bootstrap 95% confidence intervals. H) ACh signal during non-rewarded trials in Control mice in the cued versus uncued reversal learning task during the first three days of reversal. Two-sample t-test, *P=0.012. N=7 Uncued, N=8 Cued.

During reversal, however, the genotype behavioral difference was substantially reduced (**Figure 7E**), with no remaining difference in perseveration (**Figure 7F**). Consistent with the model prediction, the performance difference between control and VAChT-cKO mice was smaller in the cued task than in the original uncued task (**Figure 7G**). Moreover, the ACh rebound following non-rewarded outcomes was reduced in control mice in the cued relative to the uncued task (**Figure 7H**), as predicted if an explicit omission cue decreases the need for ACh-dependent processing of ambiguous outcome-related experience. Together, these findings support a model in which DMS ACh facilitates the formation of action–outcome states when outcome-related sensory information is weak or ambiguous.

## Discussion

Although striatal acetylcholine (ACh) has been shown to be modulated by reward-predicting cues and outcomes, how it facilitates flexible learning remains unclear. Here, we studied the role of dorsomedial striatal (DMS) ACh in reversal learning. Task-evoked ACh responses were larger on non-rewarded than rewarded trials and were further modulated by reward expectation, with the largest rebound occurring when reward was omitted after animals chose the high P_reward_ lever. Reducing ACh release using vAChTcKO mice or optogenetically blunting task-evoked ACh modulation delayed reversal learning, whereas selectively enhancing the ACh modulation on non-rewarded trials facilitated reversal learning. Computational modeling suggested that the striatal ACh signal is best explained by the model-derived probability of latent state creation during behavioral updating. The circuit-constrained model predicted that ACh would be particularly important when animals must construct a new state from an uncued reward omission, but less necessary when reward omission is explicitly marked by an external cue. Consistent with this prediction, cueing non-rewarded trials reduced reversal-learning deficits in VAChTcKO mice.

### Computational modeling supports a role for ACh in state creation

An open question in reinforcement learning is how animals construct the task-relevant state space over which learning occurs^42,47^. Standard reinforcement-learning models typically assume that the state space is predefined by the experimenter, and that dopaminergic prediction errors update the values of predefined states^17^. Yet animals must construct such states by abstracting over their experiences^60^, and these internal states often differ from those assumed by experimenters^40,61^. This raises a fundamental question: how are such states constructed in the first place?

Our algorithmic-level modeling suggests that a model-derived probability of latent-state creation tracked trial-by-trial DMS ACh dynamics during reinforcement learning. Because the animals were naïve to this task, where animals never encountered a situation in which they received no reward after pressing a lever, they likely had to construct task-relevant states from experience, in particular, during non-rewarded trials. Inspired by prior work on state splitting and latent-cause inference in extinction learning^46,47^, we formulated a reinforcement learning model in which each new experience is assigned either to an existing state with similar past experiences or to a newly created state, over which the value is learned. This model captured task behavior and, when fit only to behavior, predicted trial-by-trial ACh dynamics through its inferred probability of state creation. This state-creation probability signal explained ACh dynamics better than previously proposed model-derived variables, including several forms of uncertainty^48,49,52^.

Our model differs from belief-state inference models, which select among predefined states^40,62^, and from uncertainty-dependent learning-rate modulation models^63–66^, which adjust value updating within a fixed state space^48,67^. In reversal learning, “state” can also refer to a higher-order contingency or meta-state, for example, whether left or right is currently favored^50,52,68^. Here, we used “state” to denote an action-specific representation that links outcome-related experience to reinforcement learning, over which values are updated through reward prediction errors. Because we studied naïve animals in a one-time reversal task, the task could be solved without assuming predefined higher-order meta-states. With repeated experience, animals may acquire hierarchical contexts or meta-states that capture the broader task structure, within which ACh may also play roles^50,51^.

Our circuit-constrained modeling provides one possible implementation of this computation. In the model, ACh was driven by a corticostriatal state–outcome mismatch, reflecting a discrepancy between the outcome-implied direction of plasticity and the baseline activity balance in the selected action channel. ACh-dependent gain modulation increased the recruitment of DMS spiny neurons during the plasticity window, allowing weak or unstable cortical inputs to establish more stable, action-specific corticostriatal connectivity. This mechanism is particularly important for uncued reward omission. Reward delivery repeatedly recruited a stable cortical input pattern because it was accompanied by an explicit sensory event. Reward omission, by contrast, generated a weaker and less reproducible input. The circuit-level mismatch signal closely tracked the state-creation signal inferred by the algorithmic model, thereby linking the proposed circuit mechanism to the behavioral latent-state account.

Within this framework, a newly created state need not correspond to a discrete attractor-like representation. It can initially consist of the stabilization of a functional experience-action-outcome pathway linking cortical input, action-selective striatal activity, and outcome-dependent plasticity. Such pathways could subsequently interact with distributed, attractor-like state representations across cortico-hippocampal-basal ganglia circuits^69,70^. Although our model does not specify the cortical source of the relevant input, medial prefrontal input has been proposed to contribute to DMS state representation^41^ and DMS cholinergic interneurons are modulated by latent state^68^.

Together, our computational modeling analyses suggest that DMS ACh helps compensate for the lack of reliable experience when animals must form a new state-action representation. Because no two experiences are exactly alike, forming a state that generalizes across time requires abstraction over experience. In our experiment, the strong, explicit cue that accompanied reward delivery provided reliable sensory evidence, allowing the model to form a stable rewarded state. By contrast, the absence of an external cue during reward omission made it challenging to construct a state from this ambiguous experience. Our models and data suggest that ACh compensates for this missing sensory structure, enabling the formation of a new non-rewarded state. Once reliable states are formed, the agent or network can learn their values through dopaminergic reward prediction errors. This idea is consistent with recent work showing that optogenetic stimulation of dopamine neurons needs to be paired with an explicit auditory cue to function as a reward prediction error^71^.

### Uncertainty accounts of ACh

ACh has long been proposed to signal uncertainty, including expected uncertainty within a stable context^49^ and uncertainty over latent task meta-states in the mPFC and hippocampus^52^. However, DMS ACh dynamics in our task were better explained by latent-state creation than by expected uncertainty, unexpected uncertainty^49^, or uncertainty over a predefined meta-state^52^. Prior work on striatal circuit modeling proposed that pauses in tonically active cholinergic interneurons regulate learning rates according to population uncertainty, computed as the entropy of MSN activity^28^. However, we did not observe systematic modulation of DMS ACh pause across trials in our task (**Figure S9**). Our model instead links DMS ACh modulation to state–outcome mismatch, which closely tracked the state-creation probability inferred by the algorithmic model.

### Why is ACh not needed for the acquisition of the task?

Our computational modeling and experimental data suggest why reducing ACh release spared acquisition but delayed reversal. During acquisition, rewarded outcomes were accompanied by reliable sensory events and could therefore support stable learning about the rewarded action. By contrast, reversal required animals to update behavior after the previously rewarded action produced uncued reward omissions. In the model, these experiences favored the creation of new non-rewarded states rather than simple overwriting of the original contingency. Reduced ACh made this process less efficient, consistent with delayed reversal and increased perseveration early in reversal.

### How might ACh during a non-rewarded outcome expand the striatal population available for plasticity?

Striatal ACh has been shown to stimulate DA release at dopaminergic terminals via nicotinic and muscarinic mechanisms^72,73,44^. Thus, DA may be released in response to the strong rebound in non-rewarded trials, which then could regulate cortico-striatal plasticity via activation of postsynaptic dopamine D1 and D2 receptors^74^. However, we did not observe DA release at the time of the rebound and DA release was not changed in the vAChTcKO, neither in response to reward nor to non-rewarded outcome. Similarly, when we acutely enhanced the bi-phasic ACh response at non-rewarded outcome using optogenetics, this manipulation did not affect DA levels thus arguing against ACh regulating plasticity via regulating DA release. In this context, the vAChTcKO results were surprising to us since the same vAChTcKO mice were reported to release less DA in a Pavlovian reversal learning task^16^. One explanation for this discrepancy may be that local regulation of DA release by ACh becomes important under high-effort conditions^75^ and the anticipatory nose poking in the Pavlovian task requires more effort than the single lever press in our reversal learning task. How this difference could be explained at the neuronal level is unclear, but it may involve different degrees of nicotinic receptor desensitization on DA terminals^72,76,77^. Another explanation could be regional differences within the striatum. Here, we recorded ACh in the DMS while the local ACh modulation of DA release in^16^ and^75^ was measured in the NAc. Independent of underlying cellular mechanisms, our data do not support the hypothesis that the ACh modulation at outcome facilitates reversal learning via regulating DA release.

One classical hypothesis proposes that the ACh pause enables a time window for cortico-striatal plasticity^78,79^ which allows for learning and switching to new action policies^80^. This would be consistent with our computational model although the defining feature of the ACh modulation is not just the pause but the dip-to-peak delta. Such a rebound of ACh could promote plasticity that weakens cortico-striatal connections to D1-MSNs while strengthening connections to D2-MSNs. Muscarinic mechanisms have been observed that would support such plasticity. M4-muscarinic signaling on D1-MSNs facilitate long-term depression (LTD) and oppose long-term potentiation (LTP)^81^ while acetylcholine enhances D2-MSN excitability through M1-mediated inhibition of Kir2 channels that will enhance plasticity onto D2-MSNs^82^. However, at this point it is still unclear how the bi-phasic ACh signal exactly regulates cortico-striatal plasticity. Most likely the timing of the dip and rebound to itself, and in relation to DA release, will be important^15^. Indeed, in a value-based decision-making task, learning occurred when reward cue-induced DA release followed the dip in ACh, and not when trial start cue-induced DA release led the dip in ACh^9^. Future studies recording from striatal MSNs while measuring ACh levels will address how this expands the striatal populations available for plasticity.

### Conclusion

Our results suggest that dorsomedial striatal ACh supports behavioral flexibility by enabling animals to construct action–outcome representations from ambiguous experience, such as uncued reward omissions. By linking ACh to latent-state creation, this work provides a computational framework for understanding how cholinergic modulation of corticostriatal plasticity supports adaptive learning. This framework may also help explain why disruptions in cholinergic and corticostriatal function, as seen in disorders such as Huntington’s disease and schizophrenia, are associated with impaired flexible updating and maladaptive behavior.

## Methods

### Subjects

Homozygous floxed VAChT mice (VAChT^flx/flx^)^83^ were crossed with heterozygous Tg(Drd2-cre)44Gsat mice (D2-Cre, MMRC stock #017263-UCD) and intercrossed to generate double transgenic D2cre^+/-^-VAChT^flx/flx^ (VAChTcKO)^43^ and VAChT^flx/flx^ (control) littermates. Four to six months old male and female VAChTcKO, VAChT^flx/flx^, and Chat-IRES-Cre heterozygous mice (The Jackson Laboratory, stock #031661) were used as experimental subjects. Mice were housed at a standard 12 h light–dark cycle and were given ad libitum access to food and water (when not being food restricted for the probabilistic choice task). All experimental procedures were conducted following NIH guidelines and were approved by Institutional Animal Care and Use Committees by Columbia University and the New York State Psychiatric Institute.

### Surgical procedures

Mice were anesthetized with 1–3% vaporized isoflurane in oxygen (1 L/min) and placed in a stereotaxic apparatus. Mice were injected with AAV9-hSyn-GRAB_ACh3.0 and AAV9-hSyn-dLight1.3b (Addgene catalog numbers 121922-AAV9, 135762-AAV9) in separate hemispheres at stereotaxic bregma-based coordinates AP 1.1 mm, ML: ±1.4 mm, DV: −2.9, −3.0, −3.1 mm (150 nL/DV site). For optogenetic experiments, biosensors were mixed in a 1:1 ratio with AAV5-hSyn-FLEX-rc[ChrimsonR-tdTomato], AAV5-EF1a-DIO-eNpHR3.0-mCherry (Addgene, catalog numbers 62723-AAV5 and 26966-AAV5) or AAV5-EF1a-DIO-mCherry (UNC Vector Core) and injected at the same coordinates. Following viral injection, 400 µm diameter optic fibers (Doric) were bilaterally implanted at bregma-based coordinates AP 1.1 mm, ML: ±1.4 mm, DV: −3.0 mm and cemented into place.

### Behavior

After recovering from surgery for two weeks, mice were food-restricted to 80-90% of their baseline weight and trained to retrieve evaporated milk from a reward port in an operant box (Med Associates). After mice learned to retrieve 60 out of 60 milk rewards, they began training in a continuous reinforcement task, in which mice learned to press an extended lever to earn a reward. At the same time, mice were habituated to being tethered via patch cord to a fiber photometry recording system. After mice learned to perform 60 lever presses within a 60 min period, we recorded fiber photometry signals from mice during the continuous reinforcement task for an additional 3 days to ensure they were able to perform an operant behavior while tethered and to confirm fluorescent biosensor signals. Note that during the continuous reinforcement task, each lever press was 100% rewarded; thus, before the probabilistic reversal learning task, mice had not experienced reward omission following a lever press.

We then recorded from mice over 5 days as they underwent initial learning of the probabilistic task (Acquisition phase). During each session, there were 80 trials: 40 choice trials where mice chose between two extended levers and 40 forced trials where mice could only press a single extended lever. Each lever had a different reward probability (80% or 20%), counterbalanced across mice. After 5 days of this initial learning, the reward probability of the levers switched (Reversal phase), and mice underwent similar sessions (40 choice trials and 40 forced trials) of reversal learning. On all trials, the extended lever(s) retracted 500 ms after lever press. On rewarded trials, a milk dipper was raised for 5s starting 500 ms after lever press. In the cued version of the probabilistic choice task, an 80 dB 8 kHz tone was presented for 5s on non-rewarded trials starting 500 ms after lever press.

### Fiber photometry

An RZ10x digital acquisition system with Synapse software (Tucker-Davis Technologies) was used to collect fiber photometry data. 30 µW of 465 nm light, amplitude modulated at 330 Hz, and 7 µW of 405 nm light, amplitude modulated at 210 Hz, were delivered through an optic fiber to stimulate ligand-dependent and ligand-independent (isosbestic) fluorescence from the biosensors. Fluorescent signals were 500-560 nm bandpass filtered with a minicube (Doric) before reaching a photosensor, and the digitized signals were lowpass filtered at 6 Hz and demodulated to obtain separate biosensor and isosbestic signals. A linear fit of the isosbestic signal to the biosensor signal was calculated to remove motion artifacts and calculate a dF/F signal. This dF/F signal was then z-scored based on the mean and standard deviation of the signal during intertrial intervals. Summary values from trial-averaged signals were calculated by finding the difference between the minimum and maximum values 1.3-3s after lever press (for GACh) or by finding the maximum value 1.7-2.3s after lever press (for dLight). For trial-by-trial analyses, GACh dip to peak height was calculated by finding the difference between the minimum value 0-2s after lever press, and the first peak with a width of at least 150 ms after that minimum. GACh dip duration was calculated by measuring the time from lever press to the peak height that the signal was negative.

### Optogenetics

For all optogenetic experiments, 595 nm wavelength light (Doric) was delivered at 2.5 mW via 400 µm diameter optical fibers. For test pulse experiments, 10 ms (for Chrimson) or 500 ms (for eNpHR) pulses were delivered after a 30s or 15s intertrial interval, respectively. For the tonic stimulation experiment, 10 ms pulses were delivered at 10 Hz from the onset of a probabilistic choice session to the end of the session. For eNpHR experiments, a single 500 ms pulse was delivered starting 500 ms after lever press on non-rewarded trials.

### Immunohistochemistry

At the conclusion of experiments, mice were first anesthetized with a ketamine (100 mg/kg) and xylazine (7 mg/kg) mixture then transcardially perfused with 4% paraformaldehyde (Electron Microscopy Sciences). 40 µm sections were taken in a vibratome and stored in PBS at 4 °C until they were used for immunohistochemistry. The following antibodies and dilutions were used: chicken polyclonal anti-GFP (1:1000, Abcam, ab13970), rabbit anti-DsRed (1:1000, Takara, catalog #632496), goat anti-chicken AlexaFluor488 (1:500, ThermoFisher, catalog #A-11041), and donkey anti-rabbit AlexaFluor546 (1:500, Thermo Fisher catalog #A10040).

### Quantification and statistical analyses of experimental data

Statistical analyses on experimental data (Figures 1-4,7) were performed in GraphPad Prism 11.0.0 or MATLAB 2024a. Two-tailed tests were used throughout unless otherwise specified. Summary data were tested for normality via a Shapiro-Wilk test to determine whether to use parametric or non-parametric tests. For fiber photometry and behavioral summary measures, Wilcoxon signed-rank tests or paired t tests were used to compare values. For grouped data, two-way repeated measures ANOVA or mixed-effect models were used. Bonferroni correction was used for post-hoc analyses. For all analyses, the alpha level was 0.05. Error bars and shaded error bands represent the s.e.m.

### State creation reinforcement learning model

#### Overview

We developed a reinforcement-learning algorithmic model in which latent states were created through learning rather than fixed in advance. On each trial, the agent selected an action *a*_*t*_ ∈ {left, right} and received an outcome-related experience input *x*_*t*_ ∈ ℝ^*D*^. The model inferred whether this input should be assigned to an existing latent state associated with the selected action or should recruit a new state.

The state creation process was inspired by sequential nonparametric clustering, in which each new experience is assigned to a previously formed cluster according to its similarity and prior occupancy or initiates a new cluster when existing representations provide a poor match^84,85^. This formulation was also motivated by latent-cause accounts in which animals construct task-relevant state representations by assigning experience to inferred latent causes^46^. State assignment combined a structural reuse prior with similarity between the current outcome-related input and existing state representations constructed from previous inputs. Existing states were favored for reuse according to their prior usage, whereas inputs that do not match established state representations favored state creation.

#### Outcome-related experience inputs

We modeled outcome-related experience inputs to capture a central asymmetry of the task. Reward delivery was accompanied by a relatively stereotyped, task-locked sensorimotor sequence, including extension of the automated reward dipper, its associated sound, and the animals’ approach to, and consumption from, the dipper. In contrast, reward omission was not paired with a corresponding explicit external event. Accordingly, the model treated rewarded and unrewarded outcomes as having different input statistics. The state-creation model described here, and the circuit-constrained model described below, used closely matched input architectures. These inputs are algorithmic variables used for latent-state assignment. At a broad level, they may summarize cortical representations of external and self-generated features of post-action experience, without requiring a commitment to a particular biological implementation. These input statistics were fixed a priori based on the structure of the task and were not fit to either the behavioral or neural data.

Reward delivery-related experience input was represented by a stable, sparse, non-negative vector. Let *S*_rew_ denote a fixed set of *k*_rew_ active dimensions. On rewarded trials, the input was generated as

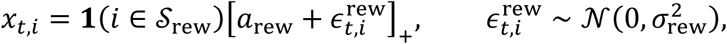

where *a*_rew_ is the input amplitude, *σ*_rew_ is the standard deviation of trial-to-trial variability, **1**(⋅) is the indicator function, and [*z*]_+_ = max (*z*, 0). The non-negative construction permitted close matching to the firing-rate-like input representation used in the circuit-constrained model. The model was initialized with one initial reward-related state for each action, reflecting the animals’ prior training with a 100% reward contingency before the reversal-learning sessions.

Because reward omission was not accompanied by a standardized external cue, we did not assign it a fixed input token. Instead, omission following each action was represented by an action-specific pattern that evolved gradually across successive trials. Let 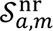 denote the set of active dimensions on the *m*-th omission experience following action *a*. On an unrewarded trial, the input was generated as

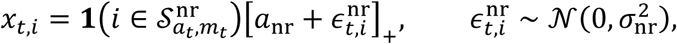

where *m*_*t*_ indexes the ordinal number of the experience following the selected action *a*_*t*_. The active set evolved independently for each action. At each trial following action *a*, each previously active dimension was retained with probability *ρ*_nr_; discarded dimensions were replaced by randomly sampled inactive dimensions, preserving *k*_nr_active dimensions. Thus, *ρ*_nr_controlled the expected overlap between successive omission-related inputs following the same action.

Input parameters were fixed rather than fit to behavioral or neural data. Unless stated otherwise, we used *D* = 20, *k*_rew_ = 3, *a*_rew_ = 2, *σ*_rew_ = 0.20, *k*_nr_ = 10, *a*_nr_ = 1, *σ*_nr_ = 0.05, and *ρ*_nr_ = 0.9.

#### Action-gated latent-state inference

Each extant latent state *j* was characterized by an outcome-related experience-input centroid ***c***_*j*,*t*_, an action label *a*_*j*_, a reward-value estimate *v*_*j*,*t*_, and a usage count *n*_*j*,*t*_. The centroid summarized outcome-related inputs previously assigned to the state, the reward-value estimate indexed its expected reward, and the usage count indexed its tendency to be reused. Although action identity could in principle be inferred from the action-specific statistics of the outcome-related inputs, we retained an explicit action label to match the action-channel structure in the circuit model. This action specificity is also natural for the behavioral task, in which outcome experiences involve distinct left and right target locations and movement from each target to the outcome location. Throughout this section, quantities with time index *t* refer to their values immediately before assignment of the outcome-related input on trial *t*; after this assignment, state variables were updated for trial *t* + 1.

For the selected action, let

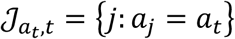

denote the set of extant states associated with the selected action and let

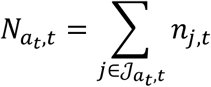

denote their total previous usage count.

#### CRP motivation and MAP-reduced implementation

The state-creation rule was motivated by the Chinese restaurant process (CRP) prior used in Bayesian latent-cause and state-discovery models^84,85^. Conditional on the selected action, the CRP assigns probability

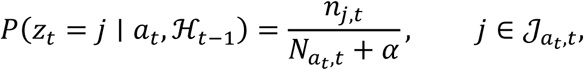

to reuse of an existing state and probability

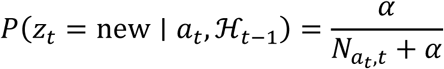

to creation of a new state. Here, *z*_*t*_ denotes the latent-state assignment of the outcome-related experience on trial *t*, ℋ_*t*−1_denotes the history available before observing ***x***_*t*_, and *α* > 0 is the CRP concentration parameter. Thus, previously used states had a greater prior probability of reuse, whereas *α* controlled the prior tendency toward state creation.

For motivation, a related generative CRP Gaussian-mixture formulation would represent each existing state by an isotropic Gaussian distribution in outcome-related input space:

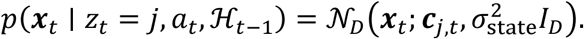

Here, ***x***_*t*_ ∈ ℝ^*D*^ is the outcome-related experience input observed on trial *t*, *N*_*D*_(***x***; ***μ***, Σ) denotes a *D*-dimensional Gaussian density with mean ***μ*** and covariance matrix Σ, ***c***_*j*,*t*_ is the centroid of state *j*, *σ*_state_ > 0 is the state-representation width, and *I*_*D*_ is the *D* × *D* identity matrix. Inputs closer to a state’s centroid were therefore more compatible with reuse of that state.

Such a fully generative model would also require a base distribution *G*_0_ over the centroid of a newly created state. The corresponding new-state prior-predictive density would be

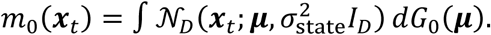

Combining the CRP prior with the Gaussian likelihood would yield unnormalized assignment weights

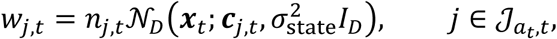

and

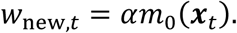

The shared denominator *N*_*at*,*t*_ + *α* cancels when these weights are normalized. This derivation shows how the CRP prior contributes a log*n*_*j*,*t*_preference for reuse of previously established states and how a new-state alternative competes with such reuse.

We did not implement this fully generative mixture model because the new-state term requires an explicit base distribution *G*_0_ and its associated parameters. Instead, the implemented model retained the CRP-derived reuse term and Gaussian similarity score for existing states, while representing the new-state alternative by an effective fitted baseline score. For each existing state,

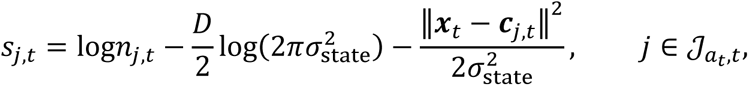

whereas the implemented new-state score was

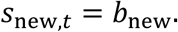

Thus, *b*_new_ is a free effective score for state creation that is a function of *α* and *m*_0_. Accordingly, *α* was introduced only in the motivating CRP formulation and was not fit separately in the implemented model.

#### MAP-reduced state-creation rule and ACh modulation

The generative CRP formulation above motivates competition between reuse of established states and creation of a new state. However, exact inference would require comparison across an expanding set of existing states and evaluation of the new-state prior-predictive density. To maintain a tractable sequential representation, we used a maximum-a-posteriori (MAP)-reduced approximation that retained only the strongest existing-state hypothesis within the selected action channel. This approximation is related to the hard-clustering approximations to Dirichlet-process Gaussian-mixture models, in which small-variance asymptotics yield a competition between assignment to the best existing cluster and creation of a new cluster^85^. This formulation is also consistent with the circuit-constrained model, with preferential recruitment of the dominant input-matching cortico-striatal representation within the selected action channel.

For the selected action, the best existing-state hypothesis was

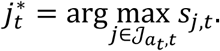

The relative log evidence favoring creation of a new state over reuse of this best existing state was

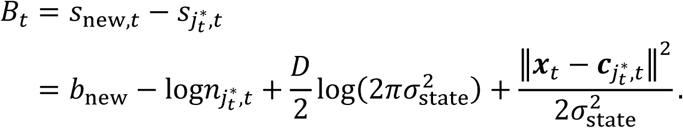

The corresponding two-alternative state-creation probability was

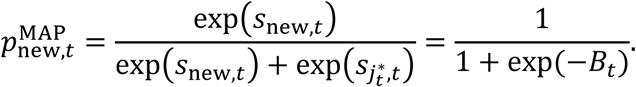

We used

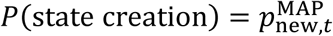

as the trialwise model-derived predictor of outcome-period ACh modulation. Thus, state creation was favored when the current input was distant from the best existing state representation or when that state had been used infrequently. If no existing state was associated with the selected action, a new state was created with probability one.

#### Binary approximation and particle filtering for behavioral fitting

For primary behavioral fitting, state assignment was approximated as binary. The outcome-related input was assigned to the more probable of the two MAP-reduced alternatives:

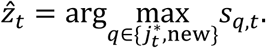

Thus, a new state was created when *B*_*t*_ > 0, equivalently when 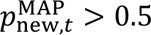; otherwise, the input was assigned to 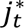. This binary approximation yielded a stable and reproducible sequential likelihood objective for parameter estimation. We validated this approximation w ith particle filtering. For particle-filter analyses and forward simulations, state assignment was sampled between the two MAP-reduced alternatives with probabilities 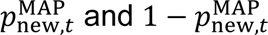, respectively (see below for implementation). To maintain the validity of the MAP reduction^85^, we constrained the state-width parameter to remain small relative to the scale of the reward-related inputs, *α*_state_ < 2 *σ*_rew_ = 0.4.

#### State updates

For binary approximation fitting, the assignment in the following updates was *z*_*t*_ = *ẑ*_*t*_. For particle-filter approximation and forward simulations, *z*_*t*_ denotes the sampled assignment. When a new state was assigned, its centroid, action label, reward value, and usage count were initialized as

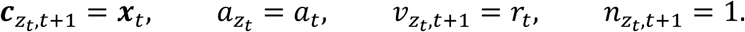

When an established state was assigned, its centroid, reward value, and usage count were updated as running averages:

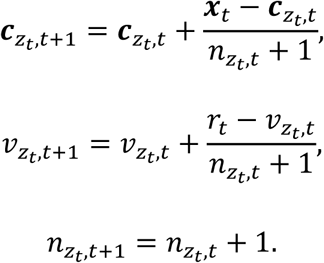

All unassigned state variables were carried forward unchanged.

#### Transition learning and action selection

The model learned action-conditioned transition probabilities over latent states. Let *T*_*t*_(*a*, *j*) denote the learned probability, before trial *t*, that action *a* leads to latent state *j*. Action values were defined as the transition-weighted expected reward value of the latent states:

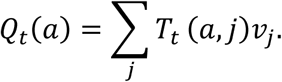

Choices were generated using a softmax policy:

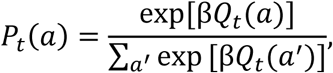

where *β* is the inverse temperature parameter.

After observing the assigned state *z*_*t*_, transition probabilities for the chosen action were updated according to

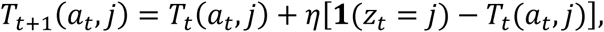

where *ή* is the transition learning rate and **1**(*z*_*t*_ = *j*) is 1 if the observed state is *j* and 0 otherwise. When a new state was created, the transition matrix was expanded to include the new state before applying this update.

#### Particle-filtering

To evaluate the stochastic MAP model, we approximated its marginal choice likelihood using a bootstrap particle filter^86^. Each of *K* particles maintained an independent realization of the evolving latent-state variables, including state centroids, usage counts, reward values, and transition probabilities.

Before trial *t*, particle *k* generated a choice probability 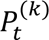 using its particle-specific state variables and the action-selection rule described above. Let 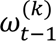 denote the normalized weight of particle *k*, with

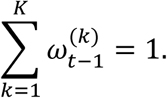

The predictive probability of the observed choice was approximated by

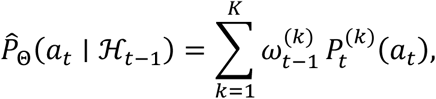

where Θ denotes the model parameters. Particle weights were updated after observing the animal’s choice,

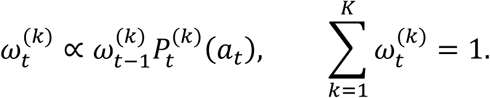

After observing the outcome, each particle updated its latent-state variables according to its sampled state assignment. The particle-filter negative log-likelihood was

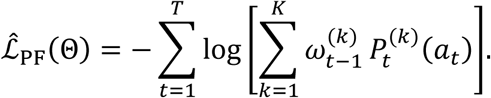

We fit the model using *K* = 30 and resampled when the effective sample size fell below *K*/2.

### Other algorithmic-level models

We tested alternative algorithmic accounts of choice behavior and trial-by-trial ACh responses: a standard Q-learning model, an uncertainty-modulated Q-learning model, and a meta-state Bayesian inference model.

#### Q-learning

We used a standard model-free Q-learning model that learned action values for the two levers^39^. The model learned the value of two actions.

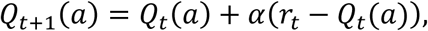

where *Q*_*t*_(*a*) is the value of action *a* on trial *t* and *α* is the learning rate. We used reward prediction error (*r*_*t*_ − *Q*_*t*_(*a*)) as a prediction of ACh modulation.

#### Uncertainty-modulated Q-learning model

The uncertainty-modulated Q-learning model was adapted from^48^. In this model, the learning rate for Q-value was modulated by expected and unexpected uncertainty. Expected uncertainty *ϵ*(*t*) was computed as a running estimate of recent unsigned reward prediction errors |*δ*(*t*)|. Unexpected uncertainty *v*(*t*) was defined as the difference between the current unsigned prediction error and expected uncertainty,

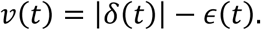

Expected uncertainty was then updated according to

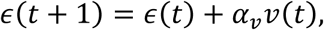

where *α*_*v*_ controls the rate of integration. Following^48^, the model used a fixed learning rate for non-negative reward prediction errors, *α*^(+)^, and a dynamic learning rate for negative reward prediction errors, 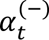. Following negative prediction errors, unexpected uncertainty modulated the subsequent negative learning rate,

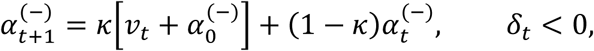

where 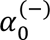 is the baseline negative learning rate and *κ* controls the rate at which unexpected uncertainty affects learning. This learning rate modulation is similar to other formulations^65^.

We used expected uncertainty, *ϵ*(*t*), and unexpected uncertainty, *v*(*t*) as trialwise ACh responses predictions.

#### Meta-state inference model

The meta-state Bayesian inference model was adapted from^52^. Briefly, the latent state, *s*_*t*_ ∈ {*L*, *R*}, indicated whether the left or right lever was currently the high-probability option. On each trial, the action–outcome pair was treated as an observation, *o*_*t*_ = {*a*_*t*_, *r*_*t*_}, where *r*_*t*_ ∈ {0,1} denotes reward omission and reward, respectively. The pre-outcome belief was *P*(*s*_*t*_ ∣ *o*^*t*−1^). The likelihood term *P*(*o*_*t*_∣ *s*_*t*_) was parameterized separately for rewarded and unrewarded trials by *c* and *d*, respectively. State transitions were governed by the parameter *γ*:

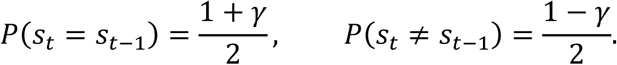

Following each observation, the belief was updated by Bayes’ rule. Following^52^, choices were generated from the pre-outcome belief using a softmax rule with fixed inverse temperature^52^ *β* = 10 and a fitted left-choice bias *b*,

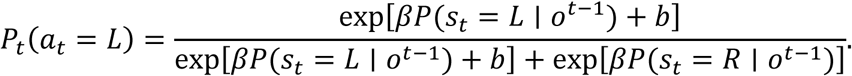

The normalized pre-outcome state uncertainty was used to predict ACh modulations^52^,

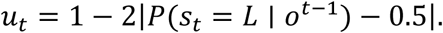

### Algorithmic model fitting, comparison, and ACh predictions

For parameter estimation and model-derived ACh analyses, each model was fit separately to each animal by minimizing the sequential negative log likelihood of observed choices using *scipy.optimize.minimize*. For standard model comparison, we used 10-fold cross-validation for each animal’s choice data and evaluated likelihood on held-out trials. To simulate behavior and ACh predictions, we generated 50 trajectories per animal using the fitted parameters. For the state-creation model, latent-state assignments were sampled according to the stochastic MAP rule described above. From these simulations, we quantified the probability of choosing the higher-reward-probability lever and the probability of staying with the same action after each action–outcome condition.

### Model robustness

To assess whether the state-creation probability signal could be reliably recovered from choice behavior, we first fit the state-creation model to each animal’s choice data. We then used the fitted animal-specific parameters to generate simulated choice datasets, with 20 simulated agents per animal. The state-creation model was then refit to each simulated dataset. We compared the trial-by-trial probability of creating a new state, P(new state), from the generative simulations with the P(new state) recovered by refitting the model to the simulated choices (**Figure S6**).

### Cortico-striatal circuit model

#### Overview

To ask how ACh shapes learning from rewarded and unrewarded outcome-related experience in the task, we built a corticostriatal circuit model in which ACh modulated outcome-phase corticostriatal recruitment and plasticity. The model incorporated standard features of dorsal striatal learning: opponent D1- and D2-MSN pathways, with D1-MSNs promoting and D2-MSNs suppressing action selection, and pathway-specific corticostriatal plasticity, with positive teaching signals favoring D1 plasticity and negative teaching signals favoring D2 plasticity in the chosen action channel. Within this framework, ACh acted as a gain signal during the outcome phase, increasing the recruitment of outcome-responsive MSNs into the plasticity window when the current corticostriatal state was poorly aligned with the obtained outcome.

The model was intended as an effective circuit-level implementation rather than a detailed biophysical mechanism. It does not require that dopamine exactly encodes the model’s teaching signal, or that cholinergic interneurons explicitly compute the mismatch term. Instead, it captures the net consequence of ACh-dependent gain modulation on outcome-phase corticostriatal activity and plasticity. Comparing simulated agents with intact versus reduced cholinergic modulation allowed us to test whether this gain-control mechanism is sufficient to support reversal learning, particularly when reward omission is uncued and therefore provides a weak or unstable cortical input.

#### Architecture

The feedforward network comprised a cortical input layer of *D* units and a striatal layer organized into two action channels, corresponding to left and right actions. Each action channel contained *M* D1-MSNs and *M* D2-MSNs. Corticostriatal weights were denoted by *W*^*L*,*D*1^, *W*^*L*,*D*2^, *W*^*R*,*D*1^, and *W*^*R*,*D*2^, where the first superscript denotes the action channel and the second denotes the MSN population.

Corticostriatal connectivity was sparse and fixed. A single binary connectivity mask *C*_*ij*_ ∈ {0,1} specified whether cortical unit *j* projected to MSN *i*. Each cortical input unit projected to a fixed subset of three MSN indices. Permitted connections (*C*_*ij*_ = 1) carried distinct weights in each action channel and pathway, whereas absent connections (*C*_*ij*_ = 0) were fixed at zero. Corticostriatal weights were constrained to remain nonnegative after each update, reflecting the excitatory nature of corticostriatal projections.

Sparse connectivity allowed distinct cortical input patterns to recruit partially overlapping but distinguishable striatal ensembles. Thus, stable cortical patterns, such as those associated with reward delivery, repeatedly engaged similar MSN ensembles, whereas drifting patterns, such as those associated with uncued reward omission, engaged partially changing ensembles across trials.

#### Trial structure and cortical input patterns

Each trial consisted of a choice phase and an outcome phase. During the *choice phase*, the model received a start-cue input, 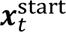, representing lever extension. Start-cue-evoked striatal activity determined action values and choice. During the *outcome phase*, the model received an outcome-related cortical input, 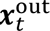, corresponding to reward delivery, uncued reward omission, or cued reward omission, depending on the simulated experiment. Outcome-evoked activity in the selected action channel then drove corticostriatal plasticity.

To link the circuit model to the state-creation reinforcement-learning model, we imposed the same qualitative input asymmetry: reward delivery was represented by a stable, reproducible input pattern, whereas uncued reward omission was represented by an action-specific pattern that changed gradually across encounters. The circuit model additionally included a start-cue input for action selection and a stable omission-related input for the cueing experiment. Numerical input amplitudes were expressed in circuit units and were set jointly with network gain, threshold, and weight parameters.

Cortical inputs were *D*-dimensional nonnegative vectors. For input type *v*, active dimension *i* was generated as

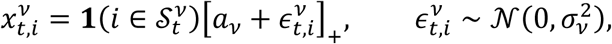

with inactive dimensions set to zero. Here, 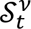 denotes the active dimensions of input type *v* on trial *t*, *a*_*v*_ is the input amplitude, *α*_*v*_ is the standard deviation of trial-to-trial variability.

The start cue and reward delivery were each represented by fixed sparse active sets, *S*_start_ and *S*_rew_, with *k*_start_ and *k*_rew_ active dimensions, respectively. Thus, reward delivery repeatedly recruited a similar cortical pattern, subject only to trial-to-trial amplitude noise.

In contrast, uncued reward omission was represented by an action-specific active set that evolved across successive omission experiences. Let 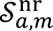 denote the active dimensions on the *m* th omission following action *a*. At each trial following action *a*, each previously active dimension was retained with probability *ρ*_nr_, and discarded dimensions were replaced by randomly sampled inactive dimensions, preserving *k*_nr_ active dimensions. Thus, *ρ*_nr_controlled the expected fractional overlap between successive uncued omission-related input patterns following the same action. Cued reward omission was represented by a fixed sparse active set, *S*_cnr_, with *k*_cnr_ active dimensions. This manipulation selectively restored the reproducibility of the omission-related input while preserving the unrewarded outcome. Outcome-related input dimensions were permitted to overlap with start-cue dimensions. This fixed overlap enabled outcome-driven plasticity to influence subsequent start-cue-driven action selection and provided a simple implementation of credit assignment.

#### Striatal activity and action selection

For action channel *c* ∈ {*L*, *R*} and pathway *p* ∈ {*D*1, *D*2}, the activity evoked by cortical input ***x*** at gain *g* was

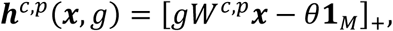

where ***h***^*c*,*p*^ is an *M*-dimensional vector of MSN activities, *θ* is a common firing threshold, **1**_*M*_ is an *M*-dimensional vector of ones, and rectification is applied element-wise.

During the choice phase, the responses to the start cue were

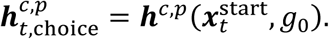

Action values combined opponent contributions from the two pathways:

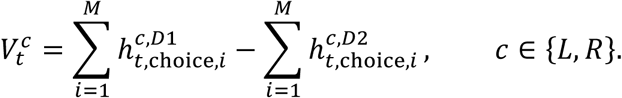

The chosen action *c*_*t*_ was sampled from a softmax policy,

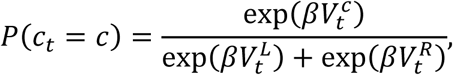

where *β* is the inverse temperature.

#### Corticostriatal state–outcome mismatch and ACh-dependent gain modulation

We modeled ACh as an outcome-phase gain signal driven by corticostriatal state–outcome mismatch. This mismatch quantified whether the baseline activity state of the selected action channel was aligned with the opponent-pathway direction of plasticity implied by the obtained outcome. Reward delivery favors a D1-dominant update, whereas reward omission favors a D2-dominant update. ACh-dependent gain was recruited when the baseline corticostriatal state opposed this outcome-appropriate update direction.

For each trial, we first computed outcome-evoked MSN activity in the selected action channel using the baseline gain, *g*_*t*_ = *g*_0_:

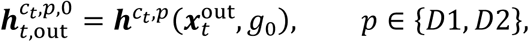

where the superscript 0 denotes baseline-gain activity. We defined a common set of outcome-responsive MSN indices,

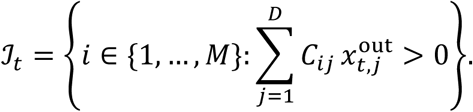

Thus, ℐ_*t*_ identified MSN indices that received input from at least one active outcome-related cortical dimension; the corresponding units were evaluated separately in each action channel and pathway through their distinct weight matrices.

We computed local baseline D1- and D2-MSN mean activities:

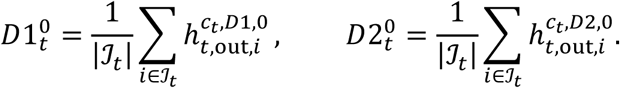

The baseline local opponent balance was

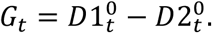

The obtained outcome specified the opponent-pathway direction of the appropriate update:

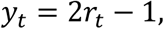

such that *y*_*t*_ = +1 for reward and *y*_*t*_ = −1 for reward omission. Outcome-congruent opponent alignment was

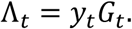

This quantity is positive when baseline local activity is aligned with the update implied by the outcome: D1 dominance after reward delivery or D2 dominance after reward omission. We defined corticostriatal state–outcome mismatch as the negative of this alignment,

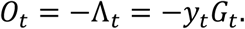

Equivalently,

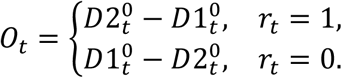

Thus, mismatch was high when D2 activity dominated after reward delivery or when D1 activity dominated after reward omission.

ACh modulation was a smooth, bounded increasing function of mismatch:

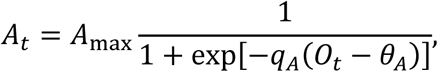

where *A*_max_ is the maximum ACh modulation, *q*_*A*_ controls the steepness of the mismatch-to-gain relationship, and *θ*_*A*_ is the mismatch threshold. The outcome-phase corticostriatal gain was then

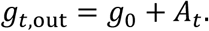

Outcome-evoked activity was recomputed using this modulated gain:

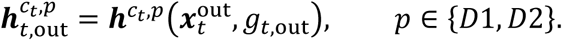

Because gain acts before the rectifying threshold, ACh modulation changed the recruitment of outcome-responsive MSNs rather than merely scaling an already active plasticity signal. In particular, increased gain could bring MSNs receiving weak outcome-related input from subthreshold to suprathreshold activity, allowing them to enter the nonzero-activity plasticity regime. This recruitment was especially consequential for weakly reproducible, drifting uncued omission inputs, whose active corticostriatal ensembles changed across encounters. In contrast, the stable cue on cued omission trials repeatedly recruited a similar ensemble and thereby reduced reliance on ACh-dependent gain. Thus, the teaching signal *δ*_*t*_ specified the direction of corticostriatal plasticity, whereas ACh-dependent gain controlled the size and composition of the outcome-responsive MSN ensemble that expressed that update.

In VAChTcKO simulations, *A*_max_ was reduced while the architecture, baseline gain, and dopamine-dependent plasticity rule were otherwise unchanged.

#### Dopamine-dependent opponent plasticity

Corticostriatal plasticity was restricted to the selected action channel. The signed teaching signal was defined from the obtained reward and the pre-outcome value of the chosen action:

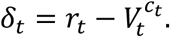

This signal has the sign convention of dopamine-dependent opponent actor plasticity: positive values favor D1 plasticity, whereas negative values favor D2 plasticity. The model assumed that this signed teaching signal remained intact across groups.

Let

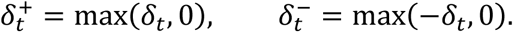

For the selected action channel, D1 weights were updated according to

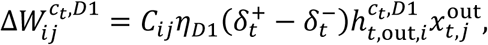

and D2 weights were updated according to

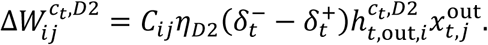

Thus, positive teaching signals strengthened active D1 synapses and weakened active D2 synapses, whereas negative teaching signals strengthened active D2 synapses and weakened active D1 synapses. The fixed connectivity mask ensured that absent projections remained absent.

Weights in the unchosen action channel were not updated on that trial. After each update, corticostriatal weights were constrained to remain nonnegative:

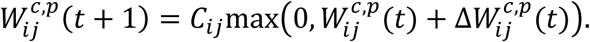

This constraint reflects the excitatory nature of corticostriatal projections, so negative updates correspond to synaptic weakening rather than a change in synaptic sign.

#### Credit assignment

Outcome-related plasticity was assigned to the action channel that produced the choice. Weights in the unchosen action channel were frozen on that trial. Outcome-related input dimensions were permitted to overlap with start-cue dimensions, enabling outcome-driven plasticity to influence subsequent start-cue-driven action selection. This fixed overlap served as a simple implementation of credit assignment. Similar choice– outcome linkage could also be implemented through eligibility traces, consistent with the temporal window of dopamine-dependent corticostriatal plasticity^87,88^.

#### Simulation parameters

Unless otherwise specified, simulations used D = 20, M = 50, β = 3, θ = 0.04, g_0_ = 2.0, q_A_ = 12, θ_A_ = 0.1, η_D1_ = η_D2_ = 0.005 and a shared fixed connectivity mask in which each cortical input dimension was projected to three MSN indices. Initial nonzero corticostriatal weights were sampled from a half normal distribution with standard deviation 0.05 and were clipped to [0,0.8]. Input parameters were k_start_ = 10, k_rew_ = 3, k_cnr_ = 3, a_start_ = 1.0, a_rew_ = 2.0, a_cnr_ = 1.2, *σ*_*start*_ = 0.02, σ_outcome_ = 0.05, a_nr_ = 1.0, ρ_nr_ = 0.7. Control agents used A_max_ = 10 and KO agents used A_max_ = 1. We simulated 15 agents per group.

### Comparison of circuit-model ACh and state-creation probability

We fit the state-creation model to the simulated action–outcome sequence from each of 100 circuit-model agents, using one parameter set per agent. We then quantified the Pearson correlation between trial-wise mean circuit-model ACh modulation and mean model-derived 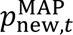 across agents.

### Permutation tests for experimental and computational-modeling analyses

Permutation tests were used to evaluate prespecified directional predictions involving measured and model-derived signals. All permutation tests were one-sided, with the direction specified by the corresponding experimental or model prediction. We used 10,000 permutations unless otherwise stated.

#### Trial-category contrasts

To test differences in behavioral, neural, or model-derived signals between trial categories during reversal, we used stratified permutation tests. Trials and sessions were treated as temporally structured components of each animal’s learning trajectory rather than as independent observations. Within each animal, trial-category labels were permuted separately within reversal-session bins, preserving the number of trials in each category within every stratum. For each permutation, the category difference was calculated within each valid stratum, averaged across strata to obtain one contrast per animal, and then averaged across animals to obtain the group-level statistic.

#### Genotype contrasts

For permutation tests involving genotype, genotype labels were permuted across animals while preserving the number of animals in each genotype group. When comparing genotype effects between the cued and uncued experiments, genotype labels were permuted separately within each experiment, thereby preserving experimental condition and group sizes. For each permutation, the control-minus-VAChTcKO contrast was calculated within each experiment, and the difference between these genotype contrasts was used as the test statistic. The observed statistic was compared with the corresponding directional tail of the permutation distribution.

### Fourier phase-randomization test for correlations between slowly varying signals

Slow temporal variations in both neural and model-derived signals can produce spurious correlations even when the signals are unrelated^59^. To address this concern, we used a Fourier phase-randomization test^58^. In this, the predicted signal was Fourier transformed, and the phase of each frequency component was randomized while preserving its amplitude spectrum. The surrogate signal was transformed back into the trial domain and correlated with the observed trial-by-trial signal. This procedure was repeated 10,000 times to construct the null distribution, preserving the signal’s autocorrelation structure.

## Author Contributions

Conceptualization and study design: D.C.L, D.Y.U, K.I., C.K. Performance of mouse experiments: D.C.L, S.O.F, M.A.M.P., S.R.E., Computational Modeling: D.Y.U., I.F.A., K.I., Data analysis: D.C.L., D.Y.U. Supervision: K.I., C.K. Manuscript preparation: D.C.L., D.Y.U., K.I., C.K. with input from all authors.

## Acknowledgements

These studies have been supported by grants from the National Institutes of Health (NIH) R01 MH124858 for C.K., R01MH136214 for K.I., T32 MH018870 and K01 MH138822 to D.C.L.

