## Supplementary Figures for "Striatal acetylcholine enables latent-state creation during reversal learning"

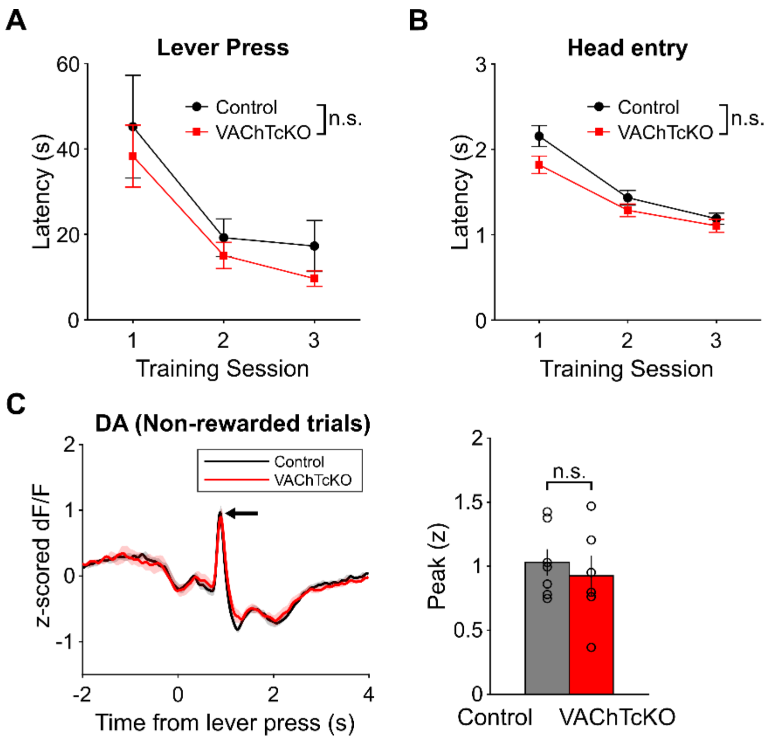


**Supplemental Figure 1: VAChTcKO does not affect DA release or operant training.**

A) Lever press latency during the first three sessions of continuous reinforcement lever press training. 2-way repeated measures ANOVA. ***P<0.0001 main effect of session, P=0.29 no main effect of genotype, P=0.96 no session*genotype interaction. N=19 Control, N=20 VAChTcKO.

B) Head entry latency during the first three sessions of continuous reinforcement lever press training. 2-way repeated measures ANOVA. ***P<0.0001 main effect of session, P=0.074 no main effect of genotype, P=0.13 no session*genotype interaction. N=19 Control, N=20 VAChTcKO.

C) Left: Comparison of DA signal on non-rewarded trials between control and VAChTcKO mice. Arrow denotes lever retraction peak. Right: Comparison of peak DA signal at lever retraction in non-rewarded trials between control and VAChTcKO mice. Two sample t-test, P=0.58. N=8 control, N=7 VAChTcKO.

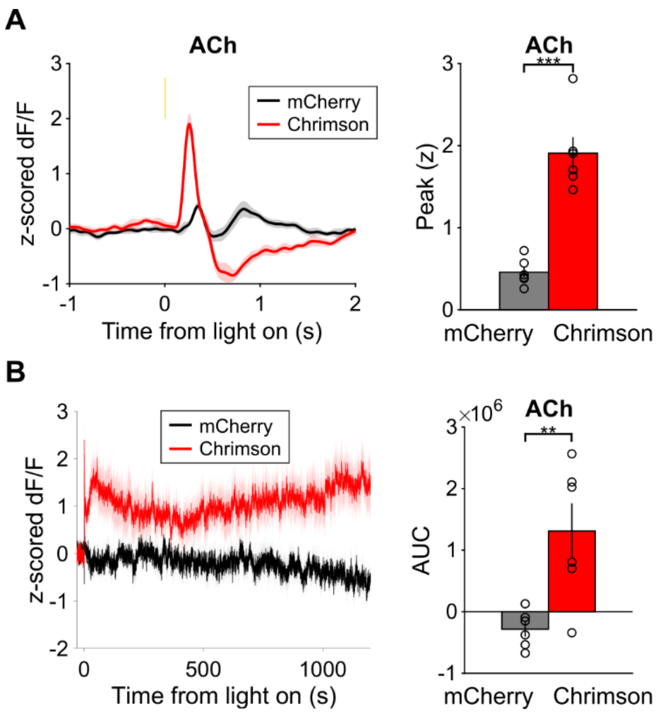


**Supplemental Figure 2: Effects of DMS CIN stimulation on ACh release.**

A) Left: DMS ACh signal in response to a single 10 ms pulse of 590 nm light for Chrimson and mCherry control mice. Bar: Time of illumination. Right: Chrimson mice exhibit a larger light-induced peak in the ACh signal compared to mCherry mice. Two sample t-test, ***P=3.6e-5. N=6 mCherry, N=6 Chrimson. Note that in naive mice under dark housing conditions light itself induces a slight ACh signal that disappears with continued exposure and under ambient light conditions of the operant box.

B) Left: ACh signals during the first 20 minutes of 10 Hz 590nm light stimulation for Chrimson and mCherry mice. Right: Area under the curve (AUC) for the first 20 minutes of light stimulation. Chrimson mice exhibit a larger AUC than mCherry mice. Two sample t-test, **P=0.0067. N=6 mCherry, N=6 Chrimson.

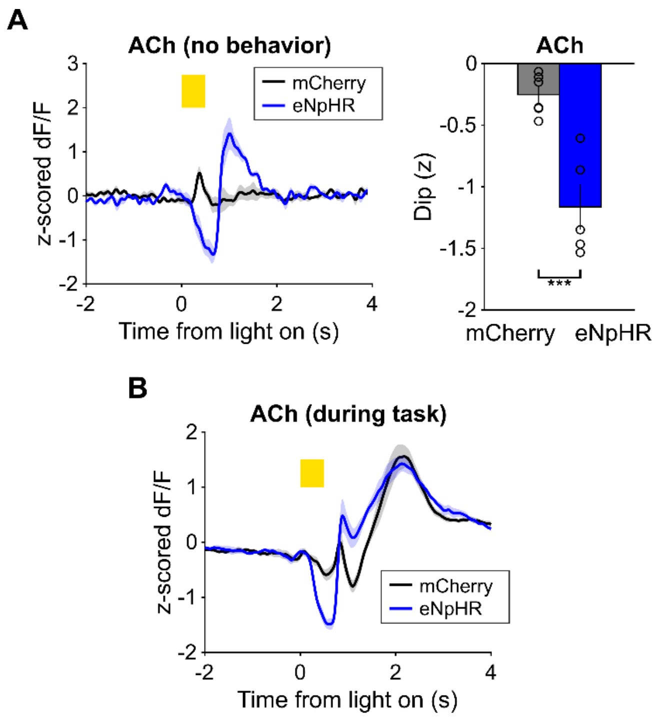


**Supplemental Figure 3: Effects of DMS CIN inhibition on ACh release.**

A) Left: DMS ACh signal in response to a single 500 ms pulse of 590 nm light for eNpHR and mCherry control mice. Bar: Time of illumination. Right: eNpHR mice exhibit a larger light-induced dip in the ACh signal compared to mCherry mice. Two sample t-test, ***P=6.9e-4. N=6 mCherry, N=5 eNpHR. Note that in naïve mice under dark housing conditions light itself induces a slight ACh signal that disappears with continued exposure and under ambient light conditions of the operant box.

B) Comparison of DMS ACh signal between eNpHR and mCherry control mice during the probabilistic reversal learning task with early light illumination. Bar: Time of illumination.

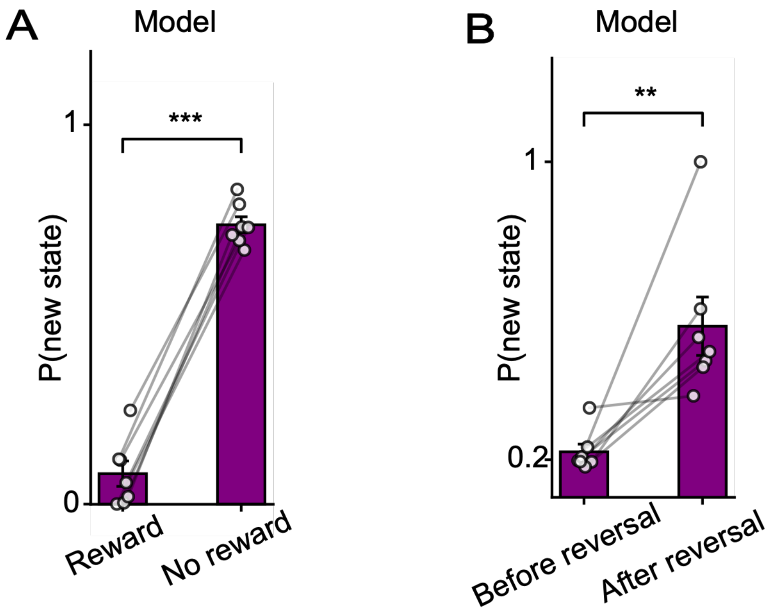


**Supplemental figure 4**: **Model-derived probability of state creation inferred from choice data**

A) Model-derived probability of creating a new state P(new state) on rewarded and unrewarded trials. The model was fit to each animal's choice data. ***p<0.001, non-parametric permutation test. Error bars indicate s.e.m.

B) Model-derived P(new state) during the final session before reversal and the first session after reversal. **p<0.01, non-parametric permutation test. Error bars indicate s.e.m.

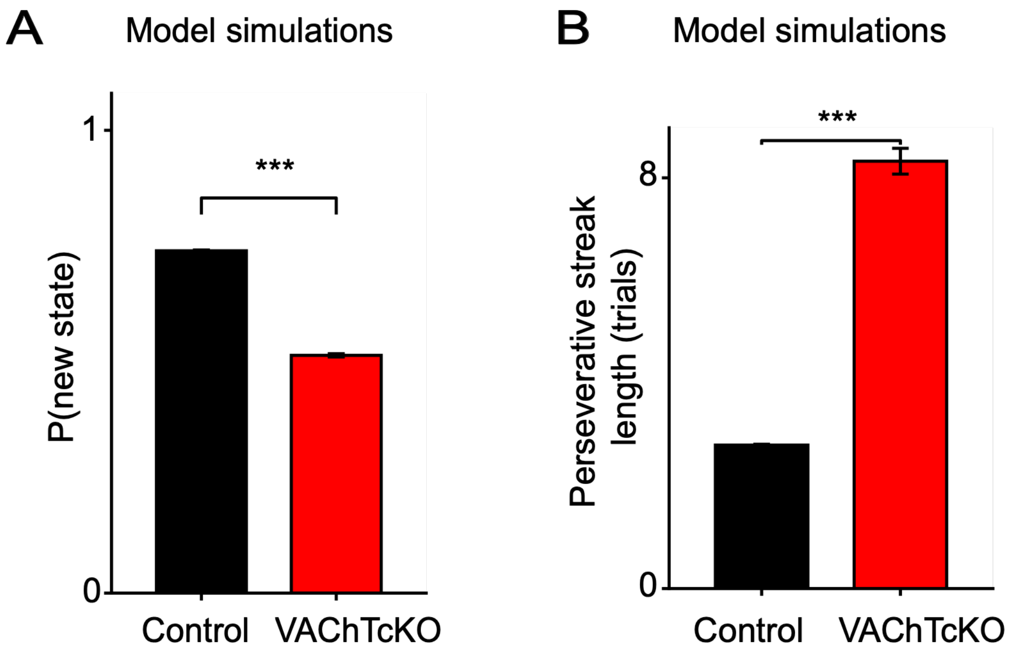


**Supplemental Figure 5: Simulations using parameters fit to control and VAChTcKO animals predict reduced state creation and increased perseveration in VAChTcKO mice.**

A) Model-derived probability of creating a new state, P(new state), during the first session after reversal in simulations using parameter sets fit to control and VAChTcKO animals. **$p<0.01$, non-parametric permutation test. Error bars indicate s.e.m.

B) Consecutive model-generated choices of the previously rewarded lever, which became the lower-reward-probability lever after reversal, during the first three reversal sessions in simulations using control and VAChTcKO parameter sets. *$p < 0.05$, non-parametric permutation test. Error bars indicate s.e.m.

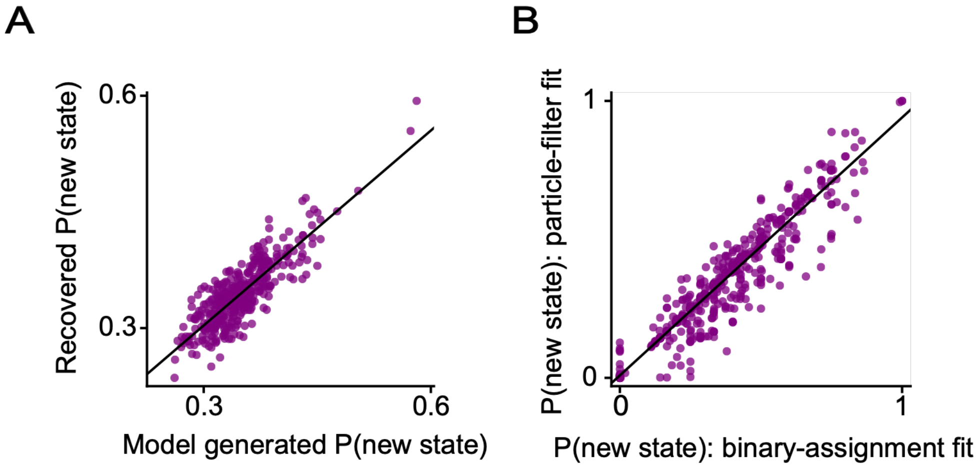


**Supplemental Figure 6: Robustness and recovery of model-derived state-creation signals.**

A) Trial-by-trial probability of creating a new state P(new state) generated during model simulations and recovered by fitting the state-creation model to the simulated choice data. Pearson correlation $r=0.85$, $p<0.0001$, Fourier phase randomization test.

B) Trial-by-trial P(new state) estimated using the binary-assignment approximation and the fitting using the particle filter. Pearson correlation $r=0.95$, $p<0.0001$, Fourier phase randomization test.

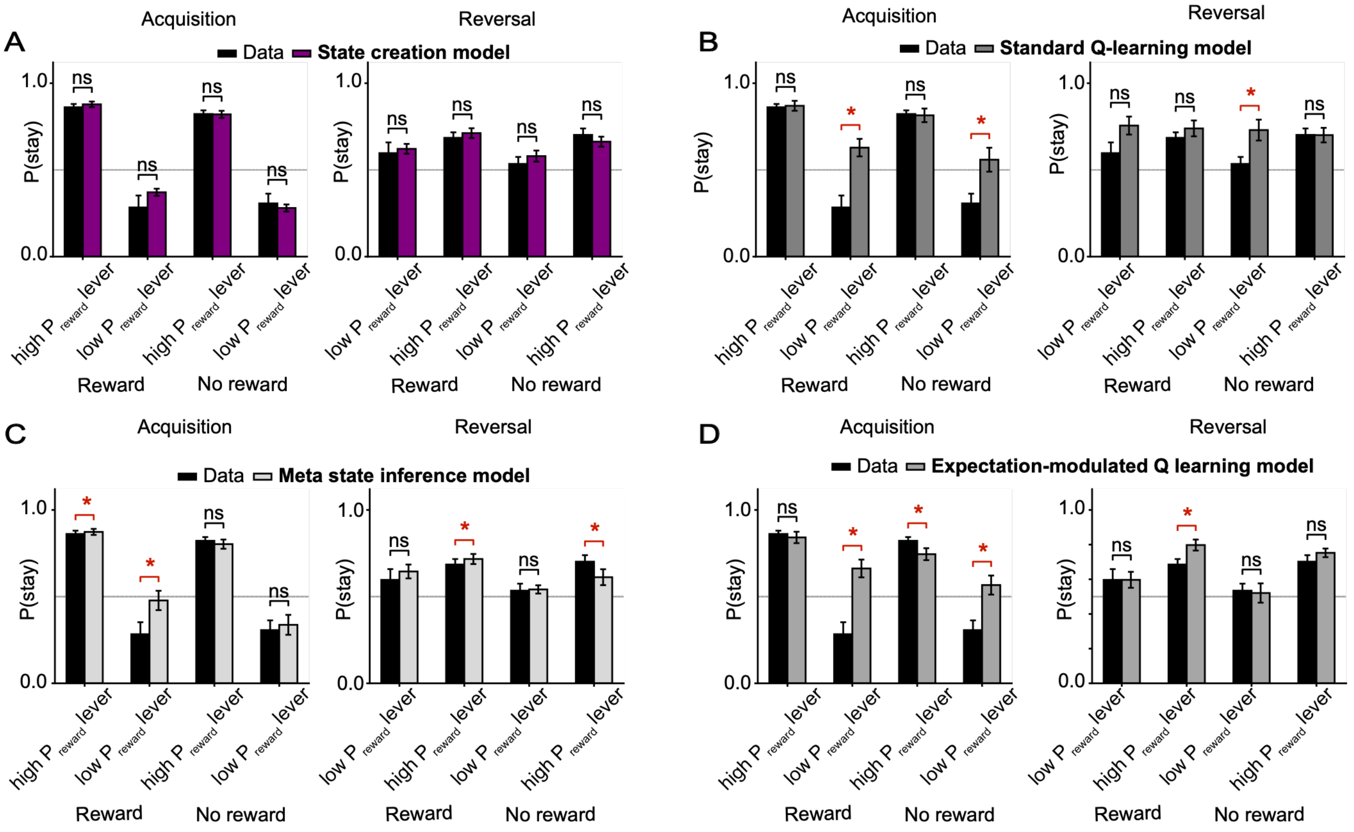


**Supplemental Figure 7: Model comparison for behavioral choice patterns.**

A) Probability of staying with the same lever on the next trial following four trial conditions: reward from the low-reward-probability lever, no reward from the low-reward-probability lever, reward from the high-reward-probability lever, and no reward from the high-reward-probability lever. Stay probabilities are shown separately for acquisition and reversal sessions in behavioral data and model simulations with the best-fit parameters.

B) Behavioral data and the state creation model.

C) Behavioral data and the Q-learning model.

D) Behavioral data and the meta-state inference model.

E) Behavioral data and the uncertainty-modulated learning rate Q-learning model.

*p<0.05, non-parametric permutation test. Error bars indicate s.e.m.

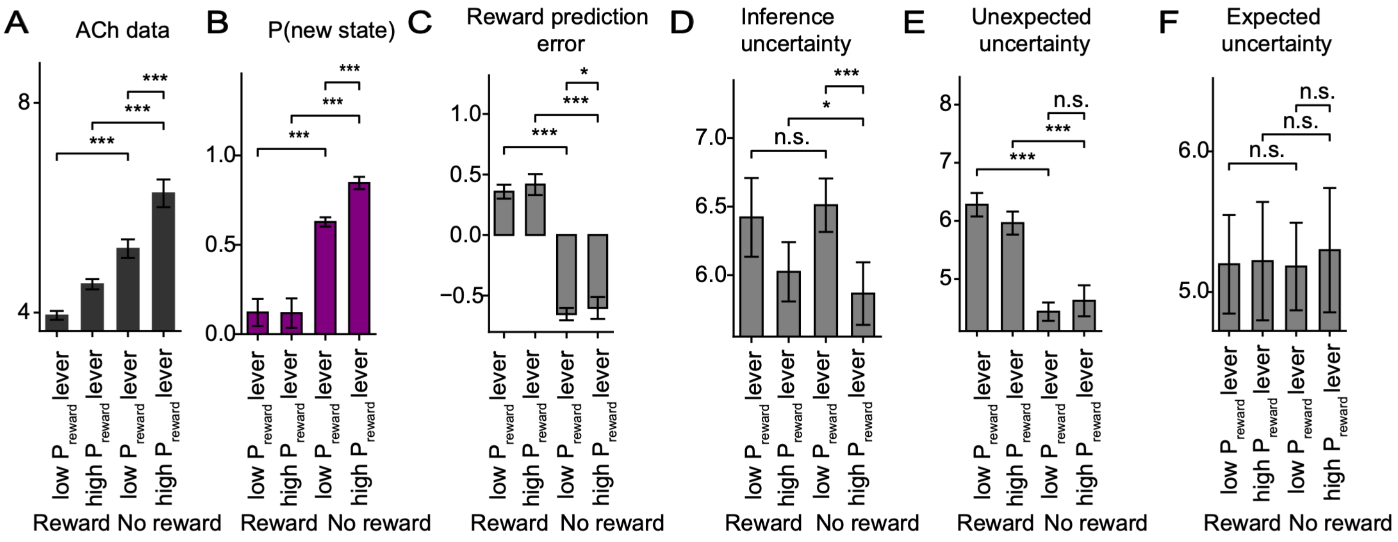


**Supplemental Figure 8: Comparison of model-derived signals with the trial-type structure of DMS ACh modulation during reversal.**

A) DMS ACh modulation across four reversal trial types: rewarded and unrewarded outcomes following choices of the high- or low-reward-probability lever.

B) State-creation model-derived probability of creating a new state P(new state), across the same trial types.

C) Reward prediction error derived from a standard Q-learning model.

D) Uncertainty derived from a meta-state inference model.

E) Unexpected uncertainty from the expectation-modulated Q-learning model.

F) Expected uncertainty from the expectation-modulated Q-learning model.

All the models were fit to each animal's choice data. $^*p<0.05$, $^{***}p<0.001$, non-parametric permutation test. Error bars indicate s.e.m.

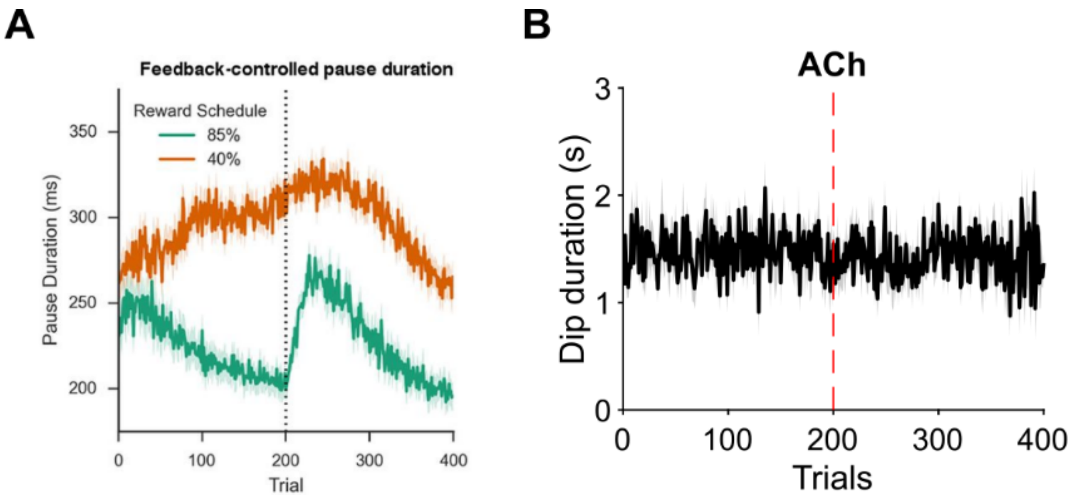


**Supplemental Figure 9: Dip duration of ACh signal does not change over learning.**

A) Figure 5 bottom left from Franklin and Frank (eLife, 2015;4 e12029). A computational model of CIN firing rate predicts that CIN pause duration will vary over the course of a 2-alternative, forced choice reversal learning task. Green trace for 85% vs 15% reward schedule, which is similar to the 80% vs 20% reward schedule used here, Orange trace for a 40% vs 10% reward schedule.

B) Dip duration of the DMS ACh signal did not change over Acquisition and Reversal. Mixed-effect model, P=0.13 no main effect of trial. N=7 mice.
